# Transcriptional and epi-transcriptional dynamics of SARS-CoV-2 during cellular infection

**DOI:** 10.1101/2020.12.22.423893

**Authors:** Jessie J.-Y. Chang, Daniel Rawlinson, Miranda E. Pitt, George Taiaroa, Josie Gleeson, Chenxi Zhou, Francesca L. Mordant, Ricardo De Paoli-Iseppi, Leon Caly, Damian F.J. Purcell, Tim P. Stinear, Sarah L. Londrigan, Michael B. Clark, Deborah A. Williamson, Kanta Subbarao, Lachlan J.M. Coin

**Affiliations:** Department of Microbiology and Immunology, University of Melbourne at The Peter Doherty Institute for Infection and Immunity, Melbourne, Victoria, 3004, Australia; Victorian Infectious Diseases Reference Laboratory, Royal Melbourne Hospital, at the Peter Doherty Institute for Infection and Immunity, Melbourne, Victoria, 3004, Australia; Department of Microbiology, Royal Melbourne Hospital, Melbourne, Victoria, 3050, Australia; Centre for Stem Cell Systems, Department of Anatomy and Neuroscience, The University of Melbourne, Melbourne, Victoria, 3010, Australia; Department of Clinical Pathology, University of Melbourne, Melbourne, Victoria, 3000, Australia; WHO Collaborating Centre for Reference and Research on Influenza, Peter Doherty Institute for Infection and Immunity, Melbourne, Victoria, 3004, Australia; Department of Infectious Disease, Imperial College London, London, Greater London, SW7 2A, United Kingdom; Institute for Molecular Bioscience, University of Queensland, Brisbane, Queensland, 4072, Australia

**Keywords:** SARS-CoV-2, COVID-19, coronavirus, nanopore, direct RNA sequencing, direct cDNA sequencing, discontinuous transcription, RNA modification, differential expression, poly(A) tail

## Abstract

SARS-CoV-2 uses subgenomic (sg)RNA to produce viral proteins for replication and immune evasion. We applied long-read RNA and cDNA sequencing to *in vitro* human and primate infection models to study transcriptional dynamics. Transcription-regulating sequence (TRS)-dependent sgRNA was upregulated earlier in infection than TRS-independent sgRNA. An abundant class of TRS-independent sgRNA consisting of a portion of ORF1ab containing *nsp1* joined to ORF10 and 3’UTR was upregulated at 48 hours post infection in human cell lines. We identified double-junction sgRNA containing both TRS-dependent and independent junctions. We found multiple sites at which the SARS-CoV-2 genome is consistently more modified than sgRNA, and that sgRNA modifications are stable across transcript clusters, host cells and time since infection. Our work highlights the dynamic nature of the SARS-CoV-2 transcriptome during its replication cycle. Our results are available via an interactive web-app at http://coinlab.mdhs.unimelb.edu.au/.

## Introduction

SARS-CoV-2, a positive-strand RNA beta-coronavirus, is the causative agent of COVID-19 (Zhou et al., 2020). As with all identified coronaviruses, the replicative and infectious cycle of SARS-CoV-2 is characterised by a process termed discontinuous minus-strand extension which occurs during replication of viral RNA by the viral Replication and Transcription Complex (RTC) within the host cell. The RTC halts synthesis of negative sense RNA when it encounters a 6 – 8 nucleotide (nt) Transcript Regulating Sequence (TRS) in the body of the genome (TRS-B) and reinitiates synthesis via a template switching event with a homologous TRS present in the 5’ leader sequence (TRS-L) (V’Kovski et al., 2020). This results in a set of nested negative-strand templates (shown in **Figure S1**) which are utilised for expression of subgenome mRNA (sgRNA). Each sgRNA includes the 3’ polyadenylated (poly(A)) Untranslated Region (UTR), a truncated set of 3’ Open Reading Frames (ORFs), and a common 5’ leader sequence. The production of subgenome transcripts alleviates pressure on the primary viral genome for protein synthesis and enables the translation of proteins at greater speed and concentration. Major SARS-CoV-2 TRS-dependent mRNAs have been previously described (Davidson et al., 2020, Kim et al., 2020, Taiaroa et al., 2020). However, the changes in the viral transcriptome and epi-transcriptome across the course of cellular infection have not yet been explored.

Long-read sequencing platforms can generate reads spanning the length of these sgRNA and are thus better suited to transcriptomic characterization of its highly nested transcriptome. One such platform is the MinION (Oxford Nanopore Technologies (ONT)), that can sequence either native RNA or cDNA directly without requirement for PCR amplification, therefore reducing PCR induced biases in estimation of expression levels (Ozsolak F et al., 2011). Furthermore, RNA modifications induce changes in ONT signal which enable exploration of the epi-transcriptome using direct RNA (dRNA) sequencing (Garalde et al., 2018, Kim et al., 2020).

In this manuscript, we carried out a comprehensive assessment of SARS-CoV-2 transcription. We generated more than 8 million long-read viral dRNA sequences and direct cDNA reads across multiple time points (2, 24 and 48 hours post infection (hpi)) with infected African green monkey (Vero) and Human (Calu-3, Caco-2) cell lines. We have developed an interactive web application for exploring the dynamic SARS-CoV-2 transcriptome at http://coinlab.mdhs.unimelb.edu.au/. Our dataset provides an expansive overview of the SARS-CoV-2 transcriptome and its changes throughout the course of infection. Our work will enable the development of new diagnostic tests for monitoring the progression of SARS-CoV-2 infectious cycle both *in vitro* and *in vivo*. This will assist in better understanding the mechanism-of-action of therapeutic agents and in monitoring the efficiency of the immune response to SARS-CoV-2 in vaccination studies.

## Results

### Infection dynamics are represented by changes in proportion of sgRNA

Viral RNA load was substantially higher in African green monkey Vero cells in comparison to Human Caco-2 and Calu-3 cell lines, reaching a maximum of 74% of all sequenced RNA at 24 hours post infection (hpi). In comparison, a maximum of 4% of all sequenced RNA mapped to SARS-CoV-2 in infected human cell lines at 48 hpi (**Figure 1a**). Even as early as 2 hpi, substantially more viral reads were detectable in Vero compared to Caco-2 and Calu-3 cells (**Figure S2**), suggesting a faster course of infection in Vero cells.

**Figure 1:**
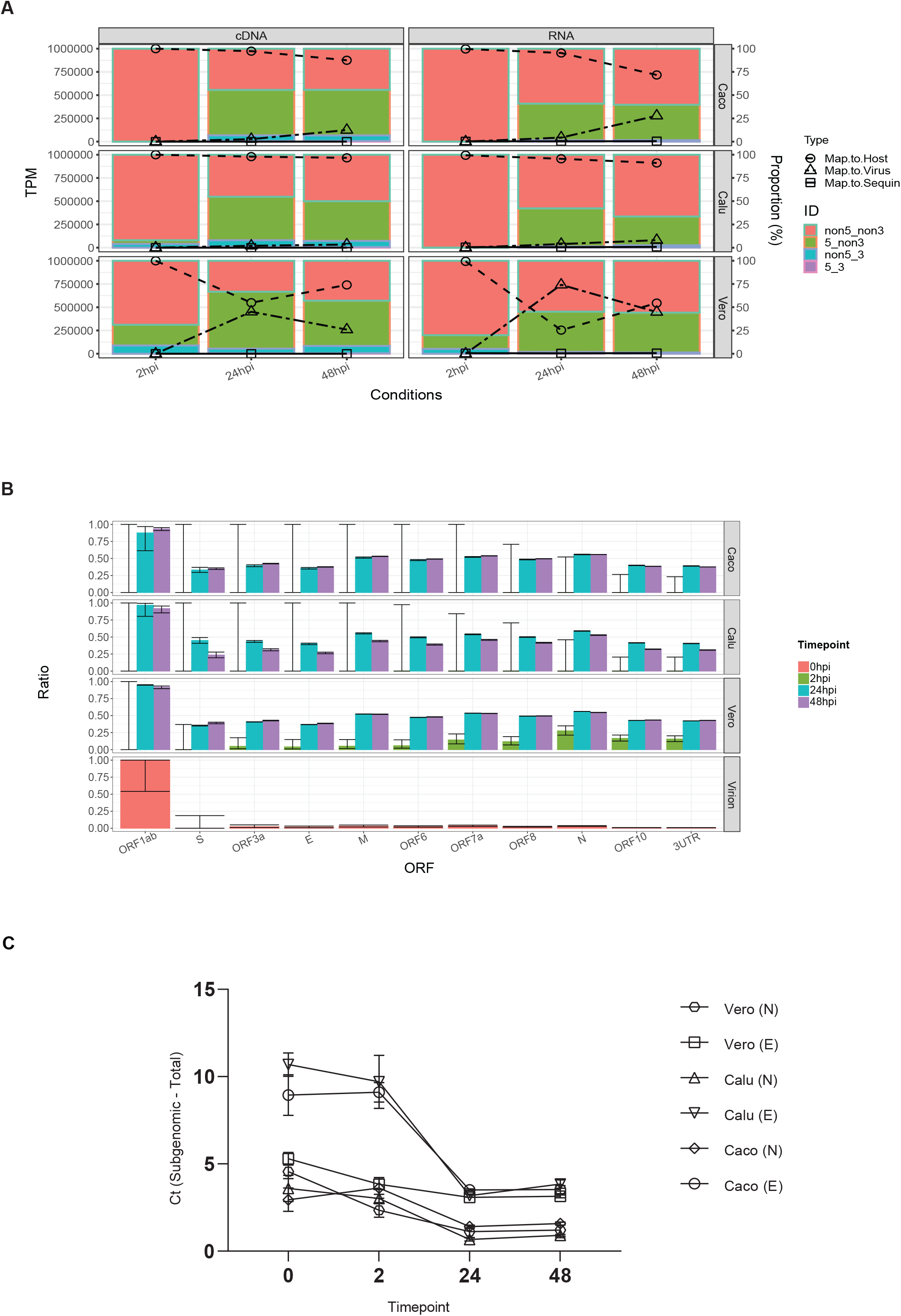
Infection dynamics are represented by changes in proportion of sgRNA. Timepoints sequenced: 2 hpi, 24 hpi and 48 hpi and cells infected: Caco, Calu, Vero. **A**. Bar chart (left axis) indicate classification of viral mapping reads based on whether they include 5’ (e.g. leader) as well as 3’ (i.e. UTR and polyA tail) in terms of transcripts per million mapped viral reads. Line graph (right axis) indicates proportion of host, viral or sequin mapping reads. **B**. Proportion of reads covering each ORF which are sgRNA by virtue of containing 5’ leader sequence for direct RNA sequencing datasets. Error bar indicates 95% CI of proportion estimate. **See also Figure S2. C**. sgRNA activity of SARS-CoV-2 measured by comparing Ct values between subgenomic and total N and E genes across all cell lines and across 4 timepoints (0, 2, 24, 48 hpi). The difference between subgenomic and total transcripts decreases over time and reaches a minimum at 24 hpi, indicating that sgRNA reaches its peak transcriptional activity at 24 hpi across all cell lines. See also **Figure S3**.

The proportion of sgRNA (i.e reads containing both 5’ leader and 3’ UTR) amongst all viral mapping reads peaked at around 40% in all three cell lines at 24 hpi (**Figure 1a**). In the non-replicating virion sample, as well as the 2 hpi samples, most reads were sequenced from the viral genome, as they had complete 3’ UTR but no 5’ leader (labelled as non5_3, **Figure S2**). This indicated that transcriptional activity had yet to accelerate at this early timepoint. Vero cells showed a greater proportion of sgRNA at 2 hpi compared to the Caco-2 and Calu-3 (Fisher exact test p=0.02)), suggesting that transcriptional activity is able to commence earlier during infection of Vero cells.

To further investigate the relationship between production of sgRNA and progression of infection, we calculated, for each ORF, the proportion of reads spanning the ORF which also contained the leader sequence (**Figure 1b**). We observed that the RNA derived from the virion sample had the least sgRNA, followed by 2 hpi, whereas the 24 hpi samples had the highest proportion of sgRNA in Vero and Calu-3 cell lines, with maximum discrimination between the virion RNA and 24 hpi obtained for the N ORF.

We then designed primers to measure both subgenomic and total N ORF expression and used quantitative Reverse Transcription PCR (RT-qPCR) with primers targeting these regions. For comparison, we applied the same approach for both subgenomic and total E ORF (Wolfel et al., 2020, Corman et al., 2020). In all three cell lines, the difference between subgenomic and total N and E ORFs was smallest at 24 hpi (1.1 and 3.1 cycle threshold (Ct) difference respectively in Vero) with a slight increase at the final 48 hpi timepoint (**Figure 1c**). This suggests that SARS-CoV-2 reaches its peak rate of transcriptional activity at the 24 hpi timepoint. By calculating expected Ct differences between subgenomic and total E and N ORFs from sequence data, we further confirmed that RT-qPCR results captured the same dynamics (**Figure S3**).

Overall, these results reveal the changing proportions of sgRNA during the SARS-CoV-2 virus infectious cycle. Our analysis using RT-qPCR to compare total and sgRNA demonstrates the potential to track viral transcriptional activity using PCR. Our data indicates that the slower rate of infection in human compared to monkey Vero cell lines may arise due to both differences in viral entry and differences in rate of early viral genome replication.

### Coronaviruses produce classes of TRS-independent sgRNA which are abundantly expressed

While all coronaviruses use a repetitive 6 nt TRS throughout the genome to generate a nested set of TRS-dependent sgRNA, the breadth of data generated in this study reveals a more detailed transcriptome that is also constituted by transcripts generated through other, unknown genome mechanisms. The depth profile of sgRNA showed sharp changes in read-depth, corresponding to negative strand disjunction mediated by TRS immediately upstream of the ORF (**Figure 2a**). To better quantify different classes of sgRNA, we developed a new tool - *npTranscript* - which assigns reads to transcript clusters (see Methods). Using *npTranscript*, we could calculate the abundance of the sgRNA at various stages of infection. At the peak of infection in Vero cells (24 hpi), the most abundant sgRNA in terms of Transcripts Per Million (TPM) mapped viral reads were ORFs N (266,000), 7a/7b (63,000), M (62,000), ORF1ab,ORF10 (60,000), ORF3a (26,000), ORF8 (16,000), ORF6 (13,000), S (7,500), E (6,100) and ORF1ab,N (5,700) (**Figure 2b**). Of these transcripts, formation of ORFs N,7a/7n, M, 3a, 8, 6, S were all mediated by a TRS-dependent homology (**Figure 2c**). The remaining two transcripts ORF1ab,ORF10 and ORF1ab,N were abundant in all 24 hpi and 48 hpi datasets and did not have breakpoints at TRS motifs (**Figure 2c**). Further inspection of TRS-independent sgRNA indicated that the majority included the first polypeptide in ORF1ab (**Figure 2d**). Taking into account polypeptide boundaries, these transcripts contained leader, *nsp1* and a variable 3’ trailer incorporating a segment of the genome upstream of ORF10 and continuing until the terminus. The exclusion of the ORF1ab stop codon will allow translation to continue into a portion of the 3’ ORF downstream of the junction site before a stop codon is reached, which has the potential to produce truncated proteins of unknown function.

**Figure 2:**
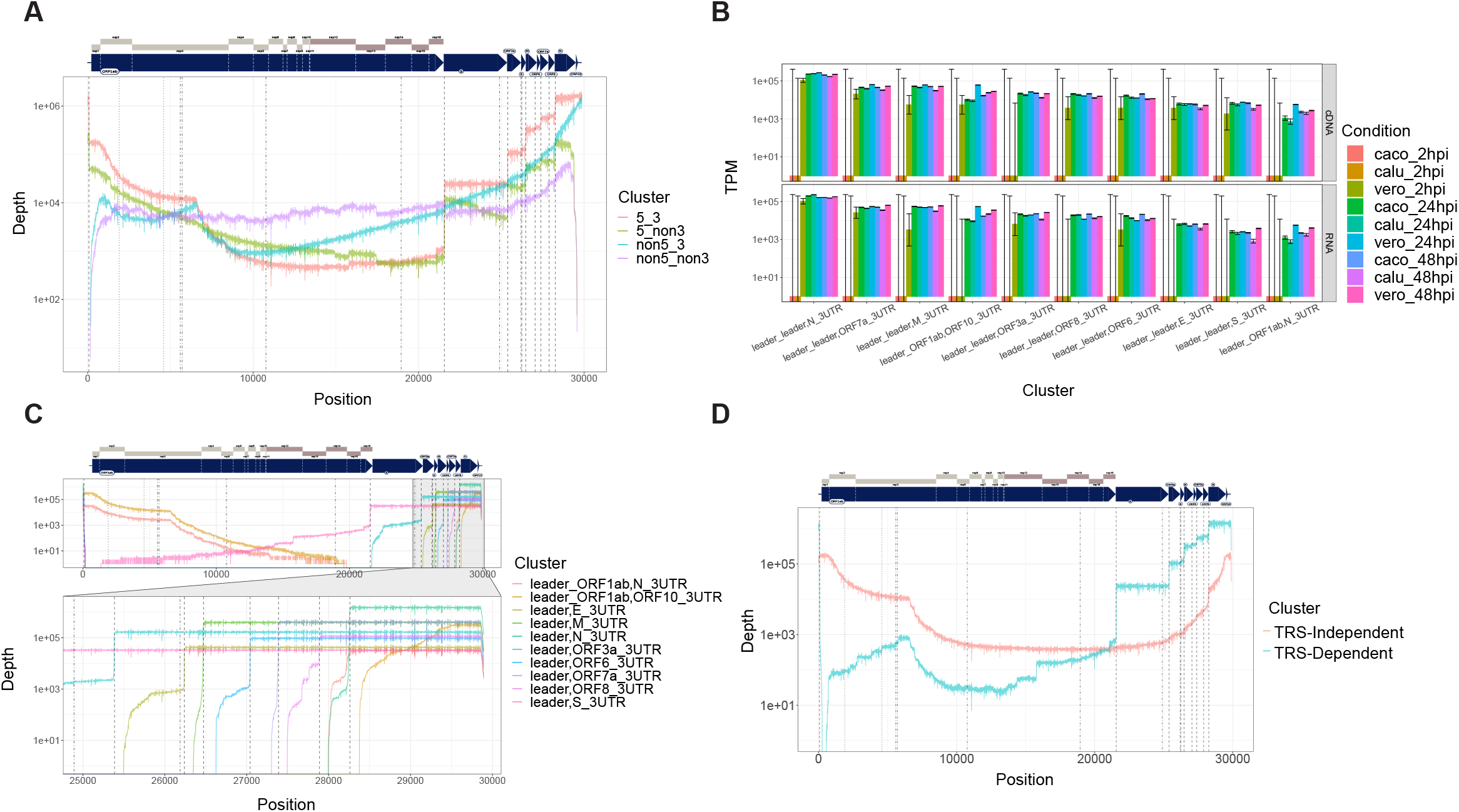
SARS-CoV-2 produces classes of TRS-independent sgRNA which are abundantly expressed. The transcript nomenclature W_X,Y_Z indicates that the transcript consists of the continuous segment from W to X joined with the segment from Y to Z. **A**. Total depth of coverage summed over all cDNA sequencing runs by categorization of read based on mapping to 5’ and 3’ end of virus (within 10 bp) plotted on a log y scale. Dashed lines indicate location of TRS motifs. **B**. Transcript abundance of major classes of sgRNA in transcripts per million mapped viral reads, plotted on a logscale for dRNA experiments (bottom row) and dcDNA (top line). 95% confidence intervals estimated from binomial model. **C**. Transcript coverage of major classes of sgRNA in terms of total read depth across all cDNA samples, shown on log scale. Dotted lines indicate positions of TRS. **D**. Coverage of TRS dependent sgRNA (blue) vs TRS independent RNA (orange), summed over all cDNA sequencing experiments. Black dashed lines indicate position of TRS motifs. Y-axis is on log scale.

To investigate whether this unusual transcript is unique to the SARS-CoV-2 (which is part of the beta-coronavirus family), we re-analysed ONT dRNA sequence data from the alpha-coronavirus 229E-HCoV (Viehweger et al., 2019). We found a similar pattern of TRS-dependent and TRS-independent sgRNA (**Figure 3a & 3c**), in which the first polypeptide nsp1 was joined to a portion of the 3’UTR. Using *npTranscript* to quantify abundance of these transcripts, we found that TRS-independent transcripts were substantially more abundant in wild-type 229E-HCoV compared to a mutant form of 229E in which the conserved 5’ Stem Loop 2 (SL2) in 229E-HCoV is replaced with that from SARS-CoV and B-CoV 3’UTR (**Figure 3b**). This finding is suggestive of a role for the SL2 of the leader sequence in the creation of these transcripts in 229E, perhaps via long-range RNA-RNA interaction, and may be relevant to the similar extended leader mRNAs found in SARS-CoV-2. Inspection of the RNA secondary structure of ORF10 + 3UTR indicates that ORF10 forms a Bulged Stem Loop (BSL) structure, upstream of the hypervariable BSL region of 3’UTR (**Figure 3d**). The BSL is a conserved feature of beta-coronavirus genomes and thought to be essential for viral replication (Madhugiri et al., 2016). Taken together, this evidence supports the role of ORF10 as part of the 3’UTR of SARS-CoV-2.

**Figure 3:**
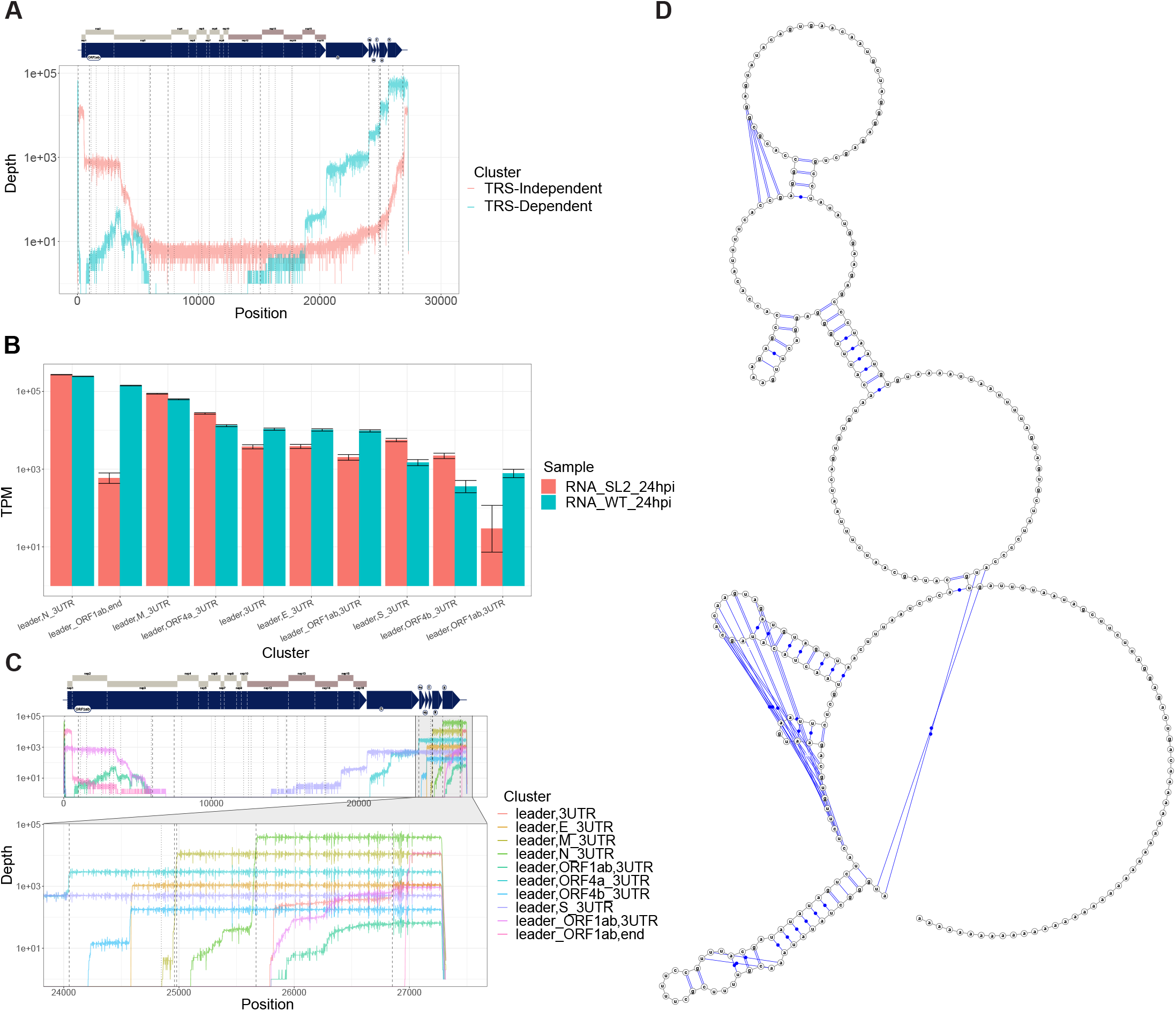
Alphacoronavirus HCoV-229E produces classes of TRS-independent sgRNA which are abundantly expressed. The transcript nomenclature W_X,Y_Z indicates that the transcript consists of the segment from W to X joined with the segment from Y to Z. **A**. Total depth of TRS-independent vs TRS-dependent sgRNA in HCoV-229E. Dashed vertical lines indicate positions of TRS motifs. **B**. Normalised transcript counts (transcript per million viral-mapped reads) of major sgRNA from 229E-HCoV for wild-type (WT) or with stem loop 2 replaced (SL2). **C**. Transcript coverage of major classes of sgRNA in terms of total read depth across all cDNA samples, shown on log scale. Dotted lines indicate positions of TRS. **D**. Predicted secondary structure of ORF10 + 3’UTR from SARS-Cov-2 calculated from IPKnot software.

### SARS-CoV-2 produces double-junction sgRNA

We also identified a persistent ‘double junction’ pattern in SARS-CoV-2 transcripts. This category featured sgRNA that showed two patterns of disjunction present at low concentrations across both dRNA and cDNA datasets (**Figure 4a**). ORF10 was the most frequently added terminal 3’ ORF in double junction sgRNA (**Figure 4b**). Most first disjunctions events were TRS-dependent, although 10% used the TRS-independent ORF1ab break point as described in previous section (**Figure S4**). In contrast, most second disjunctions were non-TRS dependent, and the 3’ breakpoint mirrored the ORF1ab,ORF10 breakpoint, suggestive of shared joining mechanism controlling this second junction that differs from TRS-mediated discontinuous minus-strand extension (**Figure S4a**). Double junction sgRNA were greatest in the Calu-3 48 hpi dataset, in which we observed leader,N,ORF10 and leader,ORF7a,ORF10 as most abundant, with 1241 and 811 TPM respectively. We also observed triple-disjunction clusters at very low levels of expression, such as ORF1ab,ORF1ab,ORF1ab,ORF10 which had an estimated 60 TPM in Calu-3 48hpi dataset. The majority of final junctions of these triple-junction reads includes the ORF10 breakpoint (**Figure S4b**).

**Figure 4:**
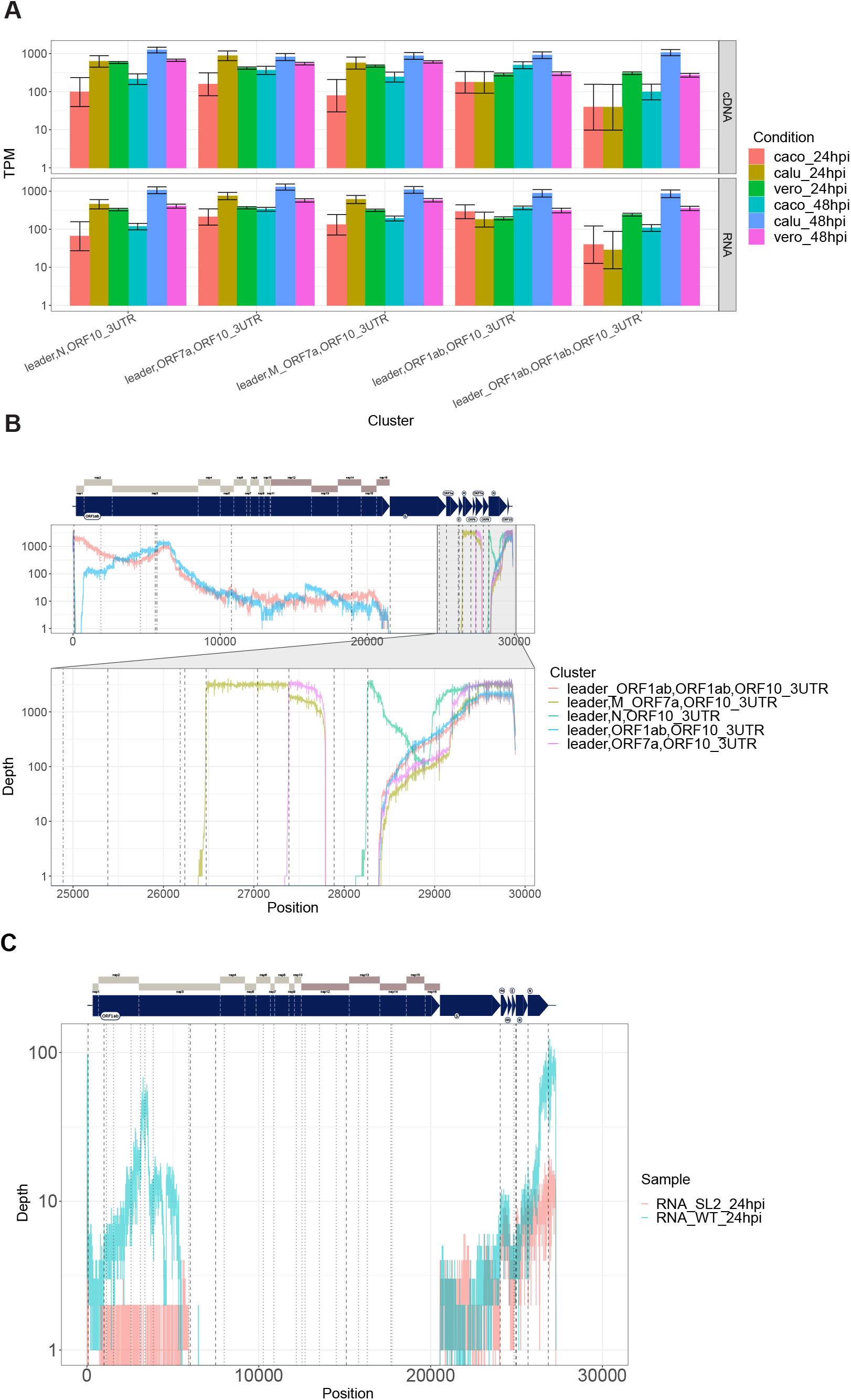
SARS-CoV-2 produces double-junction sgRNA. The transcript nomenclature U_V,W_X,Y_Z indicates that the transcript consists of the segments U to V, W to X and Y to Z. **A**. Normalised counts (in transcripts per million mapped viral reads) of double-junction reads in SARS-CoV-2 cDNA datasets. **B**. Depth of coverage of double junction sgRNA in SARS-CoV-2 (summed over all cDNA sequencing experiments), shown on log scale. Dashed lines indicate positions of TRS motifs. **C**. Coverage of double junction reads in 229E for wild-type (WT) as well as samples with modified stem loop 2. See also **Figure S4**.

In comparison, 229E-HCoV appeared to have a smaller proportion of double-junction reads. Nevertheless, we observe a leader,ORF1ab,3UTR double junction cluster (**Figure 3c**). This cluster was only observed in the WT 229E-HCoV strain, and hence highly dependent on SL2 in the 3’UTR (**Figure 4d**).

### Viral transcript polyadenylation is produced from negative strand templating rather than host factors

We detected reads with no region mapping to the 3’ end of the viral genome in both cDNA and dRNA datasets **(Figure 1a**). Upon inspection with *Nanopolish* ‘polya’ we found that no poly(A) tail is detected in most of these reads (**Figure S5a**), and that more than half of these reads also lacked detectable sequencing adapter. This observation was consistent regardless of whether the transcripts mapped to the viral 5’ terminus and contrasted with transcript categories that mapped to the 3’ end and possessed clearly segmented poly(A) tails. This observation was consistent with polyadenylation being produced by templating from the negative strand, rather than from host polyadenylation factors. The quantity of non-3’ reads varied from 2 – 4% (median = 3.3%) of viral reads in all the dRNA datasets we analysed except for Vero 24 hpi in which 10% of reads lacked the expected 3’ viral segment (**Table S1**). We found a non-random distribution of terminal breaks for non-3’ reads, however the sequence composition of their end segments does not support the idea that it is driven by runs of internal poly(A) (**Figures S5b & S5c**). Given the requirement for poly(A) tails for ONT sequencing, we considered that these reads may arise from incorrect segmentation of a single read into multiple reads, only one of which possessed a poly(A) tail.

### Viral sgRNA expression patterns change during the course of cellular infection

In order to interrogate the differential expression of SARS-CoV-2 transcriptional clusters during the time-course, we analysed ONT direct cDNA data which was sequenced in triplicate for each time point (2, 24 and 48 hpi) and each cell line (Vero, Calu-3, Caco-2). We utilized *npTranscript* to generate a reference transcriptome of sgRNA produced by SARS-CoV-2, and to assign reads to transcript clusters (see Methods), followed by *DESeq2* for differential expression analyses. We normalized each library by the number of viral mapping reads, rather than the total number of viral and host mapping reads in order to establish changes in relative abundance, rather than simply track increase in overall viral RNA during the infection (which can be seen in **Figure 1**). From this analysis, we could identify differentially expressed transcripts between timepoints – 24 vs 48 hpi (late) in all three cell lines, and in 2 vs 24 hpi (early to late) in the Vero cell line only (due to extremely low abundance of viral mapping reads in human cell lines at 2 hpi).

Interestingly, in addition to differential expression of transcripts which have both 5’ leader and 3’ UTR, we also found differential expression of transcripts which lacked the leader (non5_3) or the 3’UTR (5_non3), or both (non5_non3) (**Figures 5a-d**). For the main analysis, we proceeded to analyse the 5_3 subset of the differential expression results (**Table 1**). From our data, we estimate that the general trajectory of differential expression of SARS-CoV-2 subgenomic transcripts during an infection presents an upregulation of TRS-dependent and TRS-independent transcripts between early and late infection, and then downregulation of TRS-dependent and TRS-independent transcripts, followed by an upregulation of fragmented non5_non3 transcripts at the final stage. Of note, the transcriptional activity of TRS-independent transcripts appeared to occur faster in Vero cells compared to the human cell lines, as seen by the delayed upregulation of TRS-independent transcript in human cell lines in relation to Vero cells (**Figure 5e**).

**Table 1.** Viral sgRNA are differentially expressed between 2 vs 24 and 24 vs 48 hpi in Vero cells and 24 vs 48 hpi in Calu-3 and Caco-2 cells. The differential expression results have been filtered by p-adj < 0.05 and |log_2_FC| > 0.5 and the transcript nomenclature W_X,Y_Z indicates that the transcript consists of the segment from W to X joined with the segment from Y_Z.

| Transcript | Log<sub>2</sub>FC | p-adj |
| --- | --- | --- |
| **Vero 2 vs 24** |  |  |
| leader_leader,N_3UTR | 2.87 | 9E-37 |
| leader_leader,ORF7a_3UTR | 2.95 | 1E-10 |
| leader_ORF1ab,ORF10_3UTR | 5.04 | 1E-07 |
| leader_leader,M_3UTR | 5.03 | 1E-07 |
| leader_leader,ORF3a_3UTR | 6.23 | 7E-06 |
| leader_leader,ORF8_3UTR | 3.08 | 0.001 |
| leader_ORF1ab,N_3UTR | 4.07 | 0.001 |
| leader_leader,ORF6_3UTR | 3.31 | 0.003 |
| leader_leader,S_3UTR | 3.45 | 0.006 |
| leader_leader,E_3UTR | 2.28 | 0.035 |
| **Vero 24 vs 48** |  |  |
| leader_ORF1ab,ORF10_3UTR | -1.25 | 7E-62 |
| leader_ORF1ab,N_3UTR | -1.20 | 5E-47 |
| leader_leader,S_3UTR | -0.71 | 6E-15 |
| leader_3UTR | -1.99 | 5E-08 |
| leader_ORF1ab,3UTR_3UTR | -1.03 | 2E-04 |
| leader_ORF1ab,end | -0.92 | 0.001 |
| leader_ORF1ab,ORF3a_3UTR | -0.85 | 0.007 |
| leader_ORF1ab,ORF8_ORF8,ORF10_3UTR | -2.18 | 0.025 |
| leader_leader,ORF3a_ORF7a,end | -2.17 | 0.026 |
| leader_ORF1ab,ORF3a_ORF3a,ORF10_3UTR | -2.99 | 0.035 |
| leader_ORF1ab,S_3UTR | -1.91 | 0.042 |
| **Caco-2 24 vs 48** |  |  |
| leader_ORF1ab,ORF10_3UTR | 0.60 | 3E-06 |
| leader_ORF1ab,ORF10_N | 1.11 | 0.005 |
| leader_ORF1ab,N_3UTR | 0.74 | 0.029 |

|  | Calu-3 24 vs 48 |  |
| --- | --- | --- |
| leader_leader,M_3UTR | -1.12 | 1E-26 |
| leader_leader,N_3UTR | -1.09 | 7E-23 |
| leader_leader,ORF3a_3UTR | -1.05 | 2E-18 |
| leader_leader,ORF8_3UTR | -1.11 | 2E-16 |
| leader_leader,ORF7a_3UTR | -0.84 | 2E-15 |
| leader_leader,S_3UTR | -1.36 | 6E-12 |
| leader_leader,E_3UTR | -1.27 | 2E-10 |
| leader_ORF1ab,ORF10_3UTR | 0.81 | 2E-09 |
| leader_leader,ORF6_3UTR | -0.83 | 1E-08 |
| leader_ORF1ab,S_ORF1ab,ORF10_3UTR | 3.47 | 0.001 |
| leader_leader,S_ORF1ab,ORF10_3UTR | 1.80 | 0.025 |

**Figure 5:**
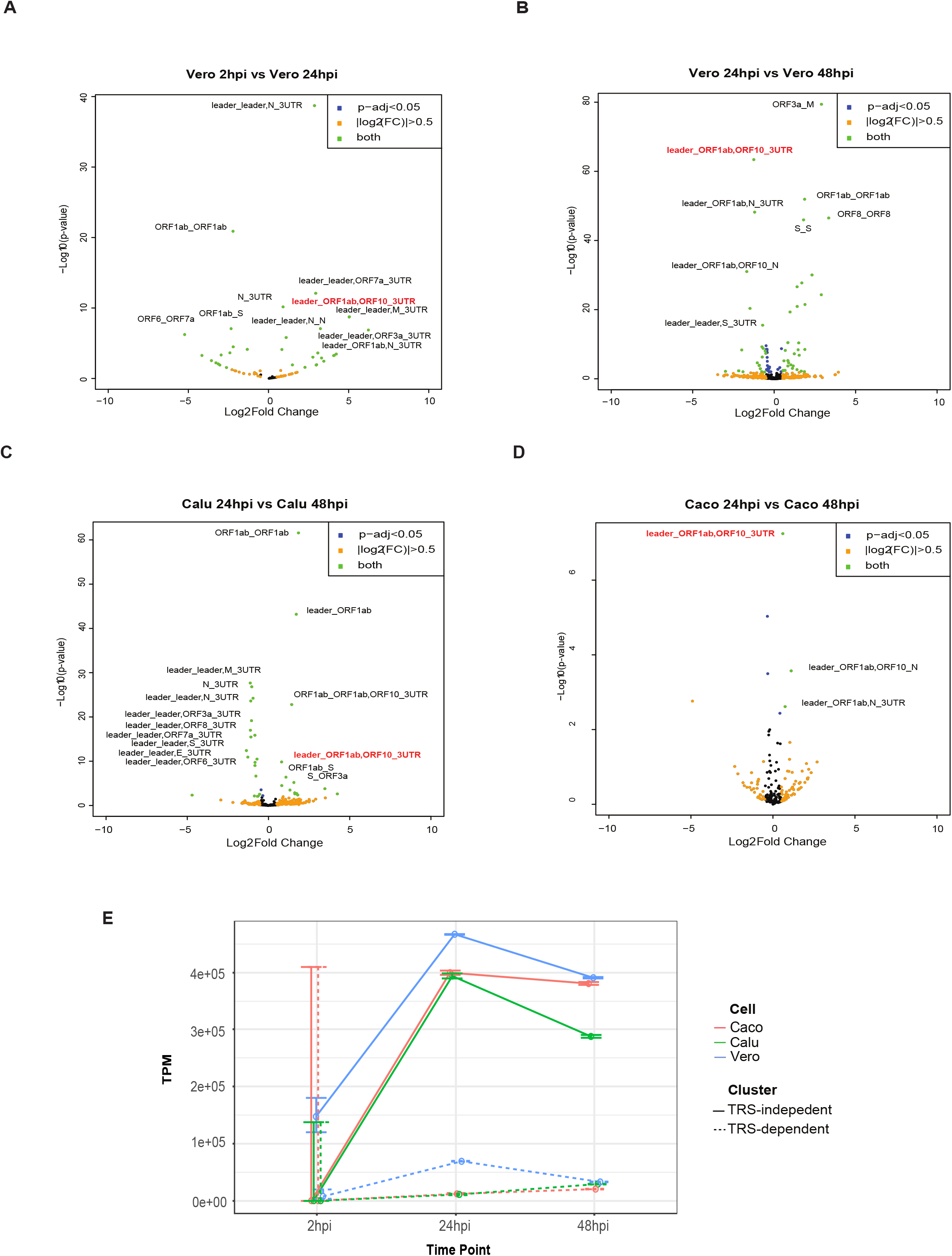
Viral sgRNA expression patterns change during the course of cellular infection with delayed responses in TRS-independent transcripts. Volcano plots of differentially expressed SARS-CoV-2 transcripts from direct cDNA datasets. **A**. Vero cells between 2 vs 24 hpi **B**. Vero, **C**. Calu-3 and **D**. Caco-2 cell lines between 24 vs 48 hpi analysed using *DESeq2*. Thresholds of p-adj < 0.05 and |log_2_FC| > 0.5 were applied to the data. Orange dots indicate transcripts which have |log_2_FC| > 0.5, blue dots indicate transcripts which have p-adj < 0.05 and green dots indicate transcripts which satisfy both criteria. Positive and negative log_2_FC indicate upregulation and downregulation at the latter timepoint, respectively. The transcript nomenclature W_X,Y_Z indicates that the transcript consists of the segment from W to X joined with the segment from Y_Z. **E**. Changes in TRS-dependent (dotted) and TRS-independent (continuous) Transcripts Per Million (TPM) mapped viral reads across multiple timepoints (2, 24, 48 hpi) in Caco-2 (orange), Calu-3 (green) and Vero (blue) cell lines. See also **Figure S8 & Table S2**.

Among these results, one TRS-independent transcript - leader_ORF1ab,ORF10_3UTR - has been shown to be consistently differentially expressed across all cell types. These transcripts were significantly upregulated (p-adj < 0.05) between 2 and 24 hpi in Vero cells, and downregulated between 24 vs 48 hpi (**Figures 5a-d**). In comparison, these transcripts are upregulated between 24 vs 48 hpi in Caco-2 and Calu-3 cells, mirroring the viral counts over time (**Figure 1a**) as the level of these transcripts peaked at 24 hpi in Vero cells and at 48 hpi in the human cell lines. Collectively, these results suggest that the peak of TRS-independent transcriptional activity occurs earlier in Vero cells compared to human cell lines and the presence of this TRS-independent transcript is of importance as they appear in all three cell types.

Additionally, we found that differentially expressed 5_3 transcripts (p-adj < 0.05) which were either genome-mapped or transcriptome-mapped revealed a positive linear correlation in log_2_FC between the two mapping methods (**Table S2**), with less transcripts being differentially expressed in transcriptome-mapped reads.

### RNA modifications vary between genomic and sgRNA but not throughout the course of infection

We used *Tombo* to determine *de novo* modification predictions on the various mRNA transcripts of the viral genome. Using virion direct RNA as baseline (Taiaroa et al., 2020), we identified changes to modification of the genome throughout the course of infection, between individual transcripts, and across the three cell lines: Vero, Calu-3, and Caco-2. The vast majority (98.2%) of reads from virion dataset included the 3’UTR but not the 5’ leader, and thus we inferred that it was almost entirely composed of reads from the viral genome rather than transcribed mRNA. The depth of coverage of this dataset was very low at the 5’ leader, and thus we are unable to report results of RNA modifications in the leader region (**Figure S6**).

The rapid infectibility of Vero cells allows a clear analysis of modifications at 24 and 48 hpi. The 2 hpi time-point failed to produce adequate subgenome expression for the analysis and only 311 viral reads were detected in total.

In our analysis, predicted viral modification sites on specific sgRNA clusters did not change markedly throughout the infection time course (**Figure 6**). However, we saw differences on sgRNA as compared to the RNA genome. In particular, all analysed sgRNA clusters displayed an absence of modification relative to virion genome in three regions as measured by the mean difference in methylated fraction (μ DMF): 26130 – 26135 in ORF3a (μ DMF = 0.62), 28858 - 28862 in ORFN (μ DMF = 0.6), & 29750**A** in 3UTR (μ DMF = 0.48) (**Figure 6**). We also observed that these modifications in the virion genome generated an artificially high rate of base-calling error at these positions (**Figure S7**).

**Figure 6.**
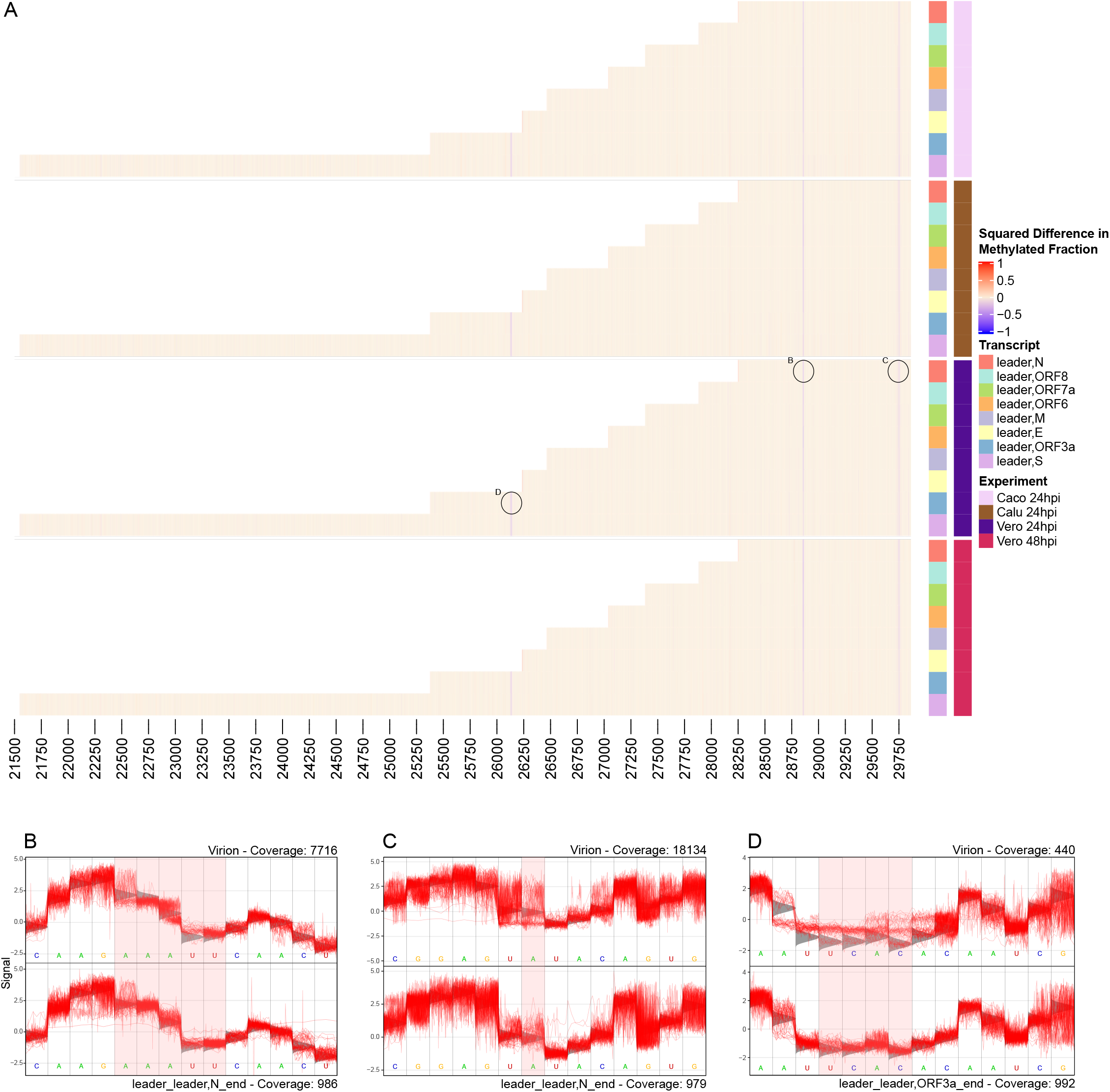
RNA modifications vary between genomic and sgRNA but not throughout the course of infection. **A**. Heatmap indicates (% age methylated reads cell line - % age methylated reads virion)^2^. sgRNA are colour-coded on the y axis and genome position is mapped on the × axis. 5’ leader sequence and final 30 bases of 3’ end are excluded due to insufficient coverage. Raw heatmap values are included in **Data S1. B, C & D**. Squiggle plots of selected significant locations from Vero 24hpi as highlighted on heatmap. Bases of interest are highlighted in red windows. Grey triangles behind squiggle indicate expected signal distribution under the standard model (i.e. no modification). For **B**, unmodified region of the N mRNA (bottom) is compared with squiggle from genomic RNA from Virion (top) signal information at genome position 28852-28862. For **C**, unmodified base at N mRNA (bottom) position 29750A compared with predicted modification on virion. For **D**, unmodified region 26310-26135 of ORF3a mRNA (bottom) compared with predicted modification on virion (top). See also **Figure S6-7 & Data S1**.

These same findings were repeated in data from Calu-3 and Caco-2 cells at 24 hpi, indicating that the different cell lines had little impact on modification changes (**Figure 6**). These results demonstrate that the viral genome carries RNA modifications which are not detectable on expressed mRNA.

The modifications reported here all consist of changes at >0.2 DMF^2^ (equivalent to DMF 0.44). A results summary including bases at >0.1 DMF^2^ (equivalent to 0.31) are included in **Data S1**.

## Discussion

The use of long-read native RNA and direct cDNA sequencing allowed the identification of TRS-dependent and independent transcripts in SARS-CoV-2. TRS-independent transcripts (sometimes referred to as non-canonical sgRNA) are formed without utilizing homologous TRS sequences and have been observed to occur in other SARS-CoV-2 transcriptome studies (Nomburg et al., 2020, Gribble et al., 2020, Kim et al., 2020, Taiaroa et al., 2020). Analysis of the time-course data presented in this manuscript shows a delayed increase of TRS-independent transcripts relative to TRS-dependent transcripts in two SARS-CoV-2-susceptible human cell lines. The most strongly upregulated of these included the leader_ORF1ab,ORF10_3UTR and leader_ORF1ab,N_3UTR transcripts (**Table 1)**.

The function of SARS-CoV-2 ORF10 remains unclear. Some studies have reported evidence of ORF10 translation (Finkel et al., 2020), while others have not found conclusive evidence of its existence in proteome databases (Taiaroa et al., 2020). Pancer et al. (2020) identified SNPs which cause premature stop-codons in ORF10 but do not impact viability *in vitro* or *in vivo*. The active transcription of the leader_ORF1ab,ORF10_3UTR transcript in our data (**Figures 5a-d**) suggests a role for ORF10 distinct from its protein coding potential. This transcript contains the full length nsp1 peptide, which is responsible for inhibiting host translation (Schubert et al., 2020), as well as the stabilizing stem loop structure from ORF10. Thus, the role of ORF10 in this context may be to stabilize the RNA molecule and enhance production of nsp1. The RNA family database RFAM (Kalvari et al., 2018) includes ORF10 in the *Sarbecovirus-3UTR*-annotated (RF03125) region of the SARS-CoV-2 genome.

We also observed ORF10 participating in the formation of the second junction in double junction transcripts. These transcripts typically have a TRS-dependent first junction, and a TRS-independent second junction to ORF10. The position of the second junction in the region upstream of ORF10 is variable, further supporting the notion that the ORF10 junction occurs in a homology independent manner.

One explanation for the mechanism of TRS-independent sgRNA formation maybe long-range RNA interactions. Long-range RNA interactions have previously been demonstrated as important for TRS-mediated leader-body joining in other coronaviruses (Mateos-Gomez et al., 2013) and may also be essential for non-TRS mediated binding as seen in SARS-CoV-2. Ziv et al. (2020) explored cis-acting RNA-RNA interactions in SARS-CoV-2 and found several long-distance interactions within ORF1ab including one which binds position 8357 nt with the 3’UTR of the genome. This interaction may be responsible for promoting generation of the ORF1ab-ORF10 transcripts we describe.

Differential expression analyses were produced by mapping to the viral genome as well as to the transcriptome (**Table S2**). Mapping to the genome allows novel transcripts to be found (Tombácz et al., 2016), whereas mapping to the transcriptome ensures the identity of the transcripts by clearly defining the junctions/breakpoints (Zhao, 2014). In this context of investigating the transcriptome of a novel coronavirus, there is more merit in mapping to the genome than the transcriptome as the transcriptome has not yet been extensively investigated and is most likely to be incomplete. The issues of using an incomplete reference transcriptome have been outlined previously (Pyrkosz et al., 2013). In our data, this is exemplified in transcripts which contain ORF1ab, as the breakpoint of ORF1ab is variable and cannot simply be defined by one breakpoint coordinate (**Figure S8**). This may explain why some transcripts are found differentially expressed in genome-mapped but not transcriptome-mapped analyses.

The data generated in this study are uniquely suited to studying the dynamics of viral epi-transcriptomics. Earlier studies have gained insights on methylation of 5’ capping for the escape of host immunity (Chen et al., 2011)(Chen et al., 2011), and the impact of host epigenetics on disease outcome (Pinto et al., 2020). Kim et al. (2020) published what is thus far the only analysis of base modifications on the viral genome in which they report 41 potential 5mC viral modification sites by contrasting signal-space information of the dRNA-sequenced viral genome against unmodified *In Vitro* Transcribed (IVT) sequence data. We used the virion-derived RNA as a control, which enabled us to focus on differences between modifications on the viral genome and transcriptome.

We find that the genomic RNA harbours more RNA modifications than the transcribed sgRNA. In particular, we report three regions which are more modified in genomic RNA than sgRNA. The strongest of these (position 28858 – 28862) was reported by Kim et al. (2020) who also reported that position 28859 is more modified amongst longer sgRNA. We extended this finding by showing that the modified state is representative of the genome RNA. We also report a remarkably stable pattern of modifications that showed very little change across cell lines and time points in transcribed sgRNA. This is the first evidence reported for the stability of SARS-CoV-2 epi-transcriptome throughout infection.

A deeper understanding of the SARS-CoV-2 transcriptome, and how it changes during infection may lead to new avenues for therapeutic strategies. One example is development of strategies to disrupt the complex patterns of negative strand disjunction to form sgRNA. Our work also highlights the importance of TRS-independent transcripts in the infectious cycle of SARS-CoV-2, which may also be an avenue for therapeutic development. Moreover, such knowledge also spurs the next generation of diagnostics for monitoring infection progression. The RNA genome modifications described here may also be a target for therapy, although further research is required to understand the role of the modifications described here.

## Data availability

The datasets supporting the results presented here are available in the NCBI repository BioProject PRJNA675370 (data currently under embargo and to be released upon publication). Transcript counts, coverage and base-calling error rates can be explored and exported via a webapp at http://coinlab.mdhs.unimelb.edu.au/.

## Acknowledgements

We would like to acknowledge Josh Lee and Uli Feltzmann for assistance with Shiny app. We would like to thank Georgia Deliyannis for assistance with culturing cell lines. LC is supported by a Career Development Fellowship from NHMRC (GNT1130084). This research was supported by NHMRC EU project grant from NHMRC GNT1195743. KS is supported by an NHMRC Investigator grant. The Melbourne WHO Collaborating Centre for Reference and Research on Influenza is supported by the Australian Government Department of Health. J J.-Y.C was supported by the Miller Foundation and the Australian government Research Training Programme (RTP) scholarship.

## Author Contributions

**Jessie J**.**-Y. Chang:** Methodology, Software, Validation, Formal Analysis, Investigation, Data Curation, Writing – Original Draft, Writing – Review & Editing, Visualisation

**Daniel Rawlinson:** Methodology, Software, Formal analysis, Data Curation, Writing – Original Draft, Writing – Review & Editing, Visualisation

**Miranda E. Pitt:** Methodology, Investigation, Validation, Supervision, Project administration, Writing – Original Draft, Writing – Review & Editing

**Josie Gleeson:** Software, Formal analysis, Data Curation, Visualisation

**George Taiaroa:** Writing – Review & Editing

**Chenxi Zhou:** Software, Formal analysis, Data Curation, Visualisation, Writing – Review & Editing

**Francesca L. Mordant:** Investigation, Methodology

**Ricardo De Paoli-Iseppi:** Investigation

**Leon Caly:** Writing – Review & Editing

**Damian F**.**J. Purcell:** Writing – Review & Editing **Deborah A. Williamson:** Writing – Review & Editing **Tim P. Stinear:** Writing – Review & Editing

**Sarah L. Londrigan:** Conceptualization, Methodology, Resources

**Mike B. Clark:** Conceptualization, Resources, Writing – Review & Editing, Supervision

**Kanta Subbarao:** Conceptualization, Methodology, Resources, Writing – Review & Editing, Supervision

**Lachlan J**.**M. Coin:** Conceptualization, Methodology, Software, Validation, Formal analysis, Investigation, Resources, Data Curation, Writing – Original Draft, Writing – Review & Editing, Visualisation, Supervision, Project administration, Funding acquisition

## Declaration of Interests

LJMC, MEP, JG, RDP and MBC have received support from Oxford Nanopore Technologies (ONT) to present their findings at scientific conferences. ONT played no role in study design, execution, analysis or publication. LJMC has received research funding from ONT unrelated to this project.

## Methods

### Cell culture

Cell lines were sourced from the American Type Culture Collection (ATCC) and included Vero (African green monkey kidney epithelial cells, ATCC CCL-81), Caco-2 (human intestinal epithelial cells, ATCC HTB-37) and Calu-3 (human lung epithelial cells, ATCC HTB-55) and maintained at 37 °C, 5% (v/v) CO2. Vero cells were cultured in Minimum Essential Media (MEM) (Media Preparation Unit, Peter Doherty Institute) supplemented with 10% Foetal Bovine Serum (FBS) (Sigma-Aldrich), 1X penicillin/streptomycin, 1X Glutamax (Gibco), and 15 mM HEPES (Gibco). Caco-2 cells were cultured in Dulbecco’s Modified Eagle Medium (DMEM) (Media Preparation Unit, Peter Doherty Institute) supplemented with 1X non-essential amino acids (Sigma-Aldrich), 20 mM HEPES, 2 mM L-glutamine, 1X Glutamax 2 μg/mL Fungizone solution, 26.6 μg/mL gentamicin, 100 IU/mL penicillin, 100 μg/mL streptomycin, and 20% FBS). Calu-3 cells were cultured in Advanced DMEM (Gibco) supplemented with 10% FBS, 100 IU/mL penicillin, 100 μg/mL streptomycin, 1X Glutamax. All cell lines were seeded in 4 × 6-well tissue-culture plates and maintained at 37°C for infection. The cell lines were tested for presence of mycoplasma using the MycoAlert Mycoplasma Detection Kit (Lonza). Virion culturing was carried out as per described previously (Taiaroa et al., 2020).

### Infection

One 6-well plate per cell line was used for each time point (0, 2, 24, 48 hpi) with triplicate wells for mock controls and infected cells. All four cell lines were infected with SARS-CoV-2/Australia/VIC01/2020 at a multiplicity of infection of 0.1 with infection inoculum composed of serum-free culture media and TPCK-treated trypsin (Worthington). The plates were incubated at 37 °C for 30 minutes. The 0-hour time point plates were removed from incubation for harvesting, and 2 mL of serum-free media + TPCK trypsin mixture was added to the plates for the remaining time points (2, 24, 48 hpi). The 2, 24 and 48 hpi plates were placed in the incubator in 37 °C, 5% CO2 until harvesting time.

### RNA extraction, DNase treatment and magnetic bead purification

The RNeasy Mini Kit (Qiagen) was used to extract the RNA using the ‘Purification of Total RNA from Animal Cells Using Spin Technology’ protocol with minor modifications. The modifications include the following; 600 μL of RLT buffer was added to the cells, and the lysates were homogenised using the Homogenizer columns (Invitrogen) as per the manufacturer’s guidelines. RNA was extracted using the RNeasy Mini Kit and treated with the DNase from the Turbo DNA-free Kit (Invitrogen) according to the manufacturer’s ‘rigorous DNase treatment’ protocol. The RNA in the supernatant was cleaned using RNAClean XP magnetic beads (Beckman Coulter) using the protocol ‘Agencourt RNAClean XP protocol 001298v001’. The magnetic beads were added to the RNA at 1.8X concentration and the final RNA was eluted in nuclease-free water.

### Oxford Nanopore Technologies library preparation and sequencing

Direct cDNA sequencing libraries were prepared using an input of 3 μg of total RNA (equivalent to approximately 150 ng of poly(A) + RNA) for Vero cell infections, and 1-2 μg of total RNA (equivalent to approximately 50 – 100 ng of poly(A) + RNA) for Caco-2 and Calu-3 cell infections per triplicate. RNA was converted to cDNA via the Direct cDNA sequencing kit (SQK-DCS109) and multiplexed using the native barcoding kit (EXP-NBD104 & 114). DRNA sequencing libraries were generated using an input of 6 μg of total RNA (2 μg per triplicate equivalent to ∼300 ng poly(A) + RNA) for Vero cell infections and 3 μg of total RNA (1 μg per triplicate equivalent to ∼150 ng poly(A) + RNA) for Caco-2 and Calu-3 cell infections via the SQK-RNA002 kit. Due to the absence of multiplexing, control or infected triplicates were pooled for dRNA sequencing per flow cell. Synthetic RNA controls (Hardwick et al., 2016) were spiked into samples at ∼10% of expected poly(A) + RNA content with Mix A used for infected samples and Mix B for uninfected controls. All libraries were sequenced with MinION R9.4.1 flow cells. Sequencing generated approximately 6 – 11 million reads for direct cDNA and roughly 1-3 million reads for dRNA sequencing. Raw data (FAST5 files) were basecalled using *Guppy* v3.5.2.

## Data analysis

### Transcript Discovery

FASTQ sequences were mapped to the reference SARS-CoV-2 genome from the first Australian case of COVID-19 (Australia/VIC01/2020, NCBI: MT007544.1) using *Minimap2* v2.11 with the splice option ‘-x splice’ engaged and ignoring TRS-dependent splice signal ‘-un’. Mapped sequences in the resulting BAM file were passed through a transcript discovery pipeline which annotates reads with information on the location of splice breakpoints relative to the viral genome. CIGAR strings are used to determine splice regions by continuous sequence of the N (not mapped) operator. Any splice traversing longer than 1000 bp of the viral genome is treated as a valid break and the genomic sites of the break are recorded in a vector such as [read_start, break1_5’, break1_3’, read_end]. We then convert this to an annotation-based string array. The read_start, read_end and 5’ breakpoint ends are converted to the first annotation which starts 5’ upstream of its position, or within 10 bp downstream of its position to allow for sequencing error. The 3’ breakpoint ends are converted to text based on the next 3’ downstream annotation, or within 10 bp 5’ upstream. This captures the fact that the disjunction sites occur immediately upstream of the target ORF. We note that this is different to the way a eukaryotic or prokaryotic gene annotation program would work. Finally, we convert the string array into a string via concatenation, with 5’ break to 3’ break concatenated using a comma to indicate the break. The end result of this procedure is an assignment of string ID, such as leader_leader,N_3UTR indicating the read starts in the leader sequence, has its first break point starting in leader and going to upstream of N, and finally ending within the 3’UTR. The code for this analysis is available at [https://github.com/lachlancoin/npTranscript].

### TRS Finding

Transcription Regulating Sequences (TRS) are required for leader-body joining during discontinuous minus-strand extension. TRS sites were located in the viral genome via a motif search using *FIMO* v5.1.1. The 6 bp segments of viral-mapping reads aligned to the TRS-dependent 5’ *ACGAAC 3’* TRS were extracted from the BAM file and transformed into a Position Weight Matrix to model variability in the sequence. The hexamer *5’ CTAAAC 3’* was used for locating TRSs in the 229E genome. The resulting PWM was converted into *meme* format using *jaspar2meme* from *MEME-suite* v5.1.1 (Bailey et al., 2009) and then used for scanning the full viral genome using *FIMO* from the same software suite.

### Methylation analysis

Signal-space FAST5 files were assessed to identify signal changes corresponding to RNA modifications using *Tombo* v1.5 (Stoiber et al., 2016). Having already been allocated a transcript cluster in *npTranscript*, read IDs from each of the 8 major subgenomes were down-sampled to 1000 reads. FAST5 reads were retrieved using the ‘fast5_fetcher_multi’ function in *SquiggleKit* (Ferguson and Smith, 2019) and resquiggled to the respective reference transcript. Transcript clusters with fewer than 1000 reads were abandoned for fear of generating an inaccurate assessment of methylation.

Resquiggled FAST5 reads were input into the ‘detect_modifications’ function using the ‘de_novo’ option which searches for any deviation from the TRS-dependent FAST5 signal. Outputs were converted to dampened_fraction wiggle files and exported for visualization and analysis in R. *ComplexHeatmap* (Gu et al., 2016) was used to produce heatmap plots of methylation data. *Tombo* ‘plot’ was used to generate squiggle plots at sites of interest. All R code for this analysis is available at [https://github.com/dn-ra/SARS-CoV-2_Mods].

### Poly(A) analysis

FASTQ passed and failed reads from dRNA sequencing were merged and indexed using *Nanopolish* v0.13.2 ‘index’ [https://github.com/jts/nanopolishusing] using the default parameters ‘-d $FAST5 -s sequencing_summary.txt $FASTQ’. The poly(A) tails of each read were estimated using the ‘polya’ function with the parameters ‘--reads $FASTQ –bam $BAM --genome $REFERENCE GENOME > combined.tsv’. A merged reference genome containing the SARS-CoV-2 Australia virus (Australia/VIC01/2020, NCBI: MT007544.1), host genome from Ensembl (release 100) and RNA sequin decoy chromosome genome (Hardwick et al., 2016) was used.

### Reverse-transcription quantitative PCR (RT-qPCR)

As a measure of infectivity, the difference between total and subgenomic transcripts which encode for the Nucleocapsid (N) and Envelope (E) genes was investigated using RT-qPCR. Barcoded cDNA from direct cDNA sequencing libraries were diluted to a concentration of ∼0.17 ng/μL and were amplified in triplicate using four sets of primers (1 μM input each primer) (Sigma-Aldrich) (**Table S3**) via the PowerUp SYBR Green Master Mix (2X) (Applied Biosystems). The amplification was carried out within the Quantstudio 7 Flex Real-Time PCR Systems (Applied Biosystems) with the standard cycling mode (50 °C, 2 mins; 95 °C, 2 mins; 50 cycles of 95 °C, 15 sec and 60 °C × 1 min). The data was analysed using the *QuantStudio Real-Time PCR Software* v1.3. As the Ct values of subgenomic E mRNA were undetectable (> 40 Ct) in 0 and 2 hpi timepoints in the human cell lines across duplicate runs, the Ct value was regarded as 40 for the purposes of measuring infectivity.

### Counts and proportions mapping to host, virus and sequin genes

*Samtools* v1.9 ‘view’ (Li et al., 2009) was used to generate the name and length of all the chromosomes in the host reference genome using the commands ‘–H $BAM | grep SQ | cut –f2-3 | sed ‘s/SN://g’ | sed ‘s/LN:?1\t/g’’. The number of host, virus and sequin reads mapping to the combined genome was counted using parameters ‘-F4 -F2048 -F256 –L $LIST_OF_CHROMS_IN_HOST.txt $BAM | wc –l’, ‘-F4 -F2048 -F256 $BAM MT007544.1 | wc –l’ and ‘-F4 -F2048 –F256 $BAM chrIS | wc –l’, respectively.

### Differential expression analysis

Passed and failed FASTQ files from direct cDNA sequencing were merged for each sequencing run and used for downstream differential expression analysis. Mapping was carried out with *Minimap2* v2.17 (Li, 2018) with the parameters ‘-ax splice –secondary=no’ to a merged reference genome containing the SARS-CoV-2 Australia virus (Australia/VIC01/2020, NCBI: MT007544.1), host genome from Ensembl (release 100) and RNA sequin decoy chromosome genome (Hardwick et al., 2016). Using the *npTranscript* pipeline, the viral reads were extracted from the BAM files to separate out reads which had primary mapping to the viral genome. The extracted viral reads were re-mapped to the viral genome using Minimap2 with the parameters ‘-ax splice –un’ as these parameters account for TRS-independent splice sites within the viral genome. During this process, *Featurecounts*-like count files were generated for differential expression analysis as *Featurecounts* (Liao et al., 2014) was unable to be used to generate suitable counts tables for the virus, perhaps due to the viral annotations being generated in-house using the *npTranscript* pipeline which are based on the ORF start position downstream of the 3’ break point instead of the breakpoint being considered as the start of the ORF. The raw counts from npTranscript were analysed using *DESeq2* v1.28.1 (Love et al., 2014) as per described below, where thresholds of |log_2_FC| > 0.5 and p-adj < 0.05 were applied. Transcript clusters with both 5’ leader and 3’ UTR sequences were retained in the results (**Table 1**). Furthermore, due to the low counts at 2 hpi, transcripts with direction of normalised counts conflicting with the direction of log_2_FC between 2 hpi and 24 hpi were regarded as false positives and flagged as being spurious.

In order to assess correlation of differential expression between genome-mapped and transcriptome-mapped transcripts, extracted viral transcripts from *npTranscript* which map to both the 5’ and 3’ ends of the full-length viral transcripts were isolated using a custom script. The new FASTQ reads were re-mapped to the viral genome using *Minimap2* with the parameters ‘-ax splice –un’ and the transcriptome with the default ‘-ax ont-map’. The same *Featurecounts*-like files were generated for genome-mapped reads as above, which were used for *DESeq2* analysis. For viral reads re-mapped with the viral 5_3 transcriptome generated by npTranscript, primary-mapped reads were isolated using *Samtools* ‘view -b -h - F 2308 $BAM > primary.bam’. *Salmon* v0.13.1 (Patro et al., 2017) was used for isoform quantification of alignments with the parameters ‘--noErrorModel –noLengthCorrection’ to obtain viral transcript counts which were input for differential transcript expression analysis in *DESeq2*. Threshold of p-adj < 0.05 were applied were applied for this analysis.

*DESeq2* was used to validate the results of differential expression. The counts from *npTranscript* and *Salmon* were input for gene and transcript level analysis respectively. Count matrices were filtered to remove very lowly expressed features (≤5 in total for each gene/transcript). Counts were normalised for sequencing depth within *DESeq2* prior to statistical analysis. Log2 fold changes and adjusted p-values (using the Benjamini-Hochberg method to correct for multiple testing) were calculated for each annotated gene or transcript and used to determine statistical significance. A regularised log transformation was subsequently performed on the normalised counts for visualisation. The PCA and volcano plots were made using the following code: https://gist.github.com/stephenturner/f60c1934405c127f09a6.

## Supplemental Information titles and legends

**Table S1: Summary table for quantities of non-3’ read amongst 24 hpi and 48 hpi timepoints**. Related to **Figure S5**.

**Table S2**: **Table of differentially expressed SARS-CoV-2 subgenomic mRNA between 24 vs 48 hpi timepoints in Vero, Caco-2 and Calu-3 cell lines from direct cDNA data**. The data shows positive linear correlation between significantly differentially expressed transcripts (p-adj < 0.05) from 5_3 genome-mapped and transcriptome-mapped transcripts (24 vs 48 hpi) via *DESeq2* analysis. Related to **Figure 5**.

**Table S3**: **List of primers used for assessing infectivity via real-time quantitative PCR**. Related to **Figure 1**.

**Figure S1**: **Schematic of major sgRNA described in this study**. sgRNA are listed in order of abundance. Most of the central part of the genome is not expressed in sgRNA but is translated directly from genomic RNA.

**Figure S2**: **Transcripts per million mapped viral reads for different read categories for RNA from infected cells 2 hpi (left axis) and total number of viral mapping reads (right axis)**. Top left cell is empty as virion cDNA was not sequenced. Related to **Figure 1**.

**Figure S3: Estimated cT difference between sgRNA and total RNA from cDNA sequence data**. Error bars indicate 95% CI estimated based on read depth. For virion E 2 hpi, only the lower 95% CI is shown. Related to **Figure 1**.

**Figure S4**: **Multiple junction transcript clusters. A**. Coverage of SARS-CoV-2 double-junction reads which have second breakpoint upstream of ORF10 (i.e. in the N ORF) vs those which do not. **B**. Coverage of SARS-CoV-2 triple junction reads which have second breakpoint upstream of ORF10 vs those which do not. Related to **Figure 4**.

**Figure S5: Characterisation of non-3’ mapping reads. A**. Poly(A) and sequencing adaptor detection from *nanopolish ‘*polya’. Groups without 3’ mapping from all experiments show consistent low detection of poly(A) tail. Vero 48 hpi has a comparatively high amount of poly(A) detection compared to other experiments which remains unexplained. **B**. Genome position at end of read for transcripts without 3’ end within final 2000 bases of viral genome. There is non-random distribution of final breaks indicating unexplained bias in the reads that pass through the nanopore for sequencing. **C**. 5-mer at end of non-3’ viral RNA reads. 5-mer is centered on final break of the transcript. Although non-random (**5B**), the sequencing of these transcripts cannot be explained by A-rich sequence at their ends. Related to **Table S1**.

**Figure S6: Coverage of SARS-CoV-2 virion dataset**. Related to **Figure 6**.

**Figure S7**: **Basecalling error rate by position for direct RNA extract from Virion (blue) and Vero at 24 hpi (orange)**. Size of dot indicates the −log10 p-value of the fisher exact test comparing basecalling error between two conditions. Error bars indicate estimate 95% confidence interval of base-calling rate estimation (estimated using binomial model). Only sites for which both conditions have a coverage of at least 1000 are shown, and at all positions reads are sub-sampled down to a depth of 1000. Dashed vertical lines indicate position of TRS. Related to **Figure 6**.

**Figure S8**: **Histogram of breakpoint positions for leader_ORF1ab**,**ORF10_3UTR transcript cluster. A**. Position at which reads assigned to cluster have a 5’ breakpoint or terminate (i.e. contribute to depth “ending”. Zoomed in region shows histogram of 5’ break points up to 2 kb. **B**. Position at which reads assigned to cluster have 3’ breakpoint or begin (i.e. contribute to depth “starting”). Zoomed in region shows histogram of 3’ breakpoints in final 1.5 kb. Dashed vertical lines show position of TRS motif. Related to **Figure 5**.

**Data S1: Supplementary data for Figure 5 modifications heatmap**. Sheet 1 contains the raw data for values in the heatmap. Sheet 2 contains a subset of sheet 1 with all values for bases with any one value > 0.1 or < −0.1. Red highlighted base numbers indicate positions that are near a transcript breakpoint and therefore in question. Units for both sheets is squared Difference in Methylated Fraction. Related to **Figure 6**.

