## Supplementary figures and tables for "Transcriptional and epi-transcriptional dynamics of SARS-CoV-2 during cellular infection"

**Table S1: Summary table for quantities of non-3' read amongst 24 hpi and 48 hpi timepoints.** Related to **Figure S5**.

| Data (dRNA) | Total reads | Total viral | Viral / total | Quant 5_no3 viral | Quant no5_no3 viral | % no3 |
|---|---|---|---|---|---|---|
| Vero 24h | 3074268 | 2270620 | 0.738588828 | 119002 | 121254 | 0.105810748 |
| Vero 48h | 1477641 | 650135 | 0.439981701 | 5228 | 19266 | 0.037675252 |
| Caco 24h | 1789012 | 76274 | 0.042634706 | 513 | 858 | 0.01797467 |
| Caco 48h | 3140226 | 862715 | 0.274730226 | 9743 | 14709 | 0.02834308 |
| Calu 24h | 2768172 | 105983 | 0.038286277 | 806 | 1286 | 0.019739015 |
| Calu 48h | 1056263 | 84297 | 0.079806828 | 974 | 2273 | 0.038518571 |

**Table S2**: **Table of differentially expressed SARS-CoV-2 subgenomic mRNA between 24 vs 48 hpi timepoints in Vero, Caco-2 and Calu-3 cell lines from direct cDNA data.** The data shows positive linear correlation between significantly differentially expressed transcripts (p-adj < 0.05) from 5_3 genome-mapped and transcriptome-mapped transcripts (24 vs 48 hpi) via *DESeq2* analysis. Related to **Figure 5**.

| **Transcript (5_3) (Calu-3)** | **$Log_2FC$ (Genome-mapped)** | **p-adj (Genome-mapped)** | **$Log_2FC$ (Transcriptome-mapped)** | **p-adj (Transcriptome-mapped)** |
|---|---|---|---|---|
| leader_leader,M_3UTR | -0.499036542 | 0.000174418 | -0.339145134 | 0.003146436 |
| leader_ORF1ab,S_ORF1ab, ORF10_3UTR | 4.589726392 | 0.000333632 | 4.689124263 | 0.024984433 |
| leader_leader,N_3UTR | -0.487392404 | 0.000363593 | -0.301293732 | 0.00198282 |
| leader_leader,S_ORF1ab,OR F10_3UTR | 2.316968538 | 0.002390042 | 4.706383312 | 0.024984433 |
| leader_leader,S_3UTR | -0.73058948 | 0.007852598 | -0.542396276 | 0.046983391 |
| **Transcript (5_3) (Caco-2)** | **$Log_2FC$ (Genome-mapped)** | **p-adj (Genome-mapped)** | **$Log_2FC$ (Transcriptome-mapped)** | **p-adj (Transcriptome-mapped)** |
| leader_leader,N_3UTR | -0.231123208 | 3.29E-06 | -0.23543418 | 3.6526E-06 |
| leader_ORF1ab,N_3UTR | 0.805493152 | 0.006221381 | 1.116735084 | 0.00881677 |
| **Transcript (5_3) (Vero)** | **$Log_2FC$ (Genome-mapped)** | **p-adj (Genome-mapped)** | **$Log_2FC$ (Transcriptome-mapped)** | **p-adj (Transcriptome-mapped)** |
| leader_ORF1ab,ORF10_3UT R | -0.778074749 | 3.49E-60 | -1.646905521 | 3.0619E-275 |
| leader_ORF1ab,N_3UTR | -0.799749914 | 5.86E-32 | -1.104385193 | 3.1755E-37 |
| leader_leader,S_3UTR | -0.490713343 | 1.76E-13 | -0.418044275 | 1.64256E-13 |
| leader_leader,ORF3a_3UTR | -0.259138102 | 4.00E-07 | -0.326730165 | 6.76954E-17 |
| leader_3UTR | -1.709982359 | 0.000468107 | 0.818593098 | 8.99277E-06 |
| leader_leader,M_3UTR | -0.173768773 | 0.001040815 | -0.356740125 | 1.02365E-32 |
| leader_leader,ORF8_3UTR | 0.187426966 | 0.007104081 | -0.133074185 | 0.008634737 |
| leader_leader,ORF3a_ORF7 a,ORF10_3UTR | 0.704433408 | 0.013883111 | 1.018348208 | 0.036112839 |
| leader_leader,E_3UTR | -0.183780458 | 0.015175927 | -0.38179938 | 1.41746E-07 |

**Table S3**: **List of primers used for assessing infectivity via real-time quantitative PCR**. Related to **Figure 1**.

| Target | Primers | Start | End | Sequence | Amplicon Size (bp) | Tm | Description |
|---|---|---|---|---|---|---|---|
| Subgenomic N | 1_nC19_L_F | 16 | 37 | CTTCCCAGGTAACAAACCAACC | 149 | 56.1 | Leader + N ORF |
| | 1_nC19_N_R | 28341 | 28362 | CCATTCTGGTTACTGCCAGTTG | | 56.1 | |
| Total N | 2_nC19_N_F | 28738 | 28759 | TGCAATCGTGCTACAACTTCCT | 137 | 56.9 | N ORF |
| | 2_nC19_N_R | 28853 | 28874 | TGCCTGGAGTTGAATTTCTTGA | | 54.6 | |
| Subgenomic E | 3_nC19_L_F | 44 | 66 | CGATCTCTTGTAGATCTGTTCTC | 167 | 52.2 | Leader + E ORF |
| | 3_nC19_E_R | 26360 | 26381 | ATATTGCAGCAGTACGCACACA | | 57.2 | |
| Total E | 3_nC19_E_F | 26251 | 26272 | TCATTCGTTTCGGAAGAGACAG | 131 | 54.7 | E ORF |
| | 3_nC19_E_R | 26360 | 26381 | ATATTGCAGCAGTACGCACACA | | 57.2 | |

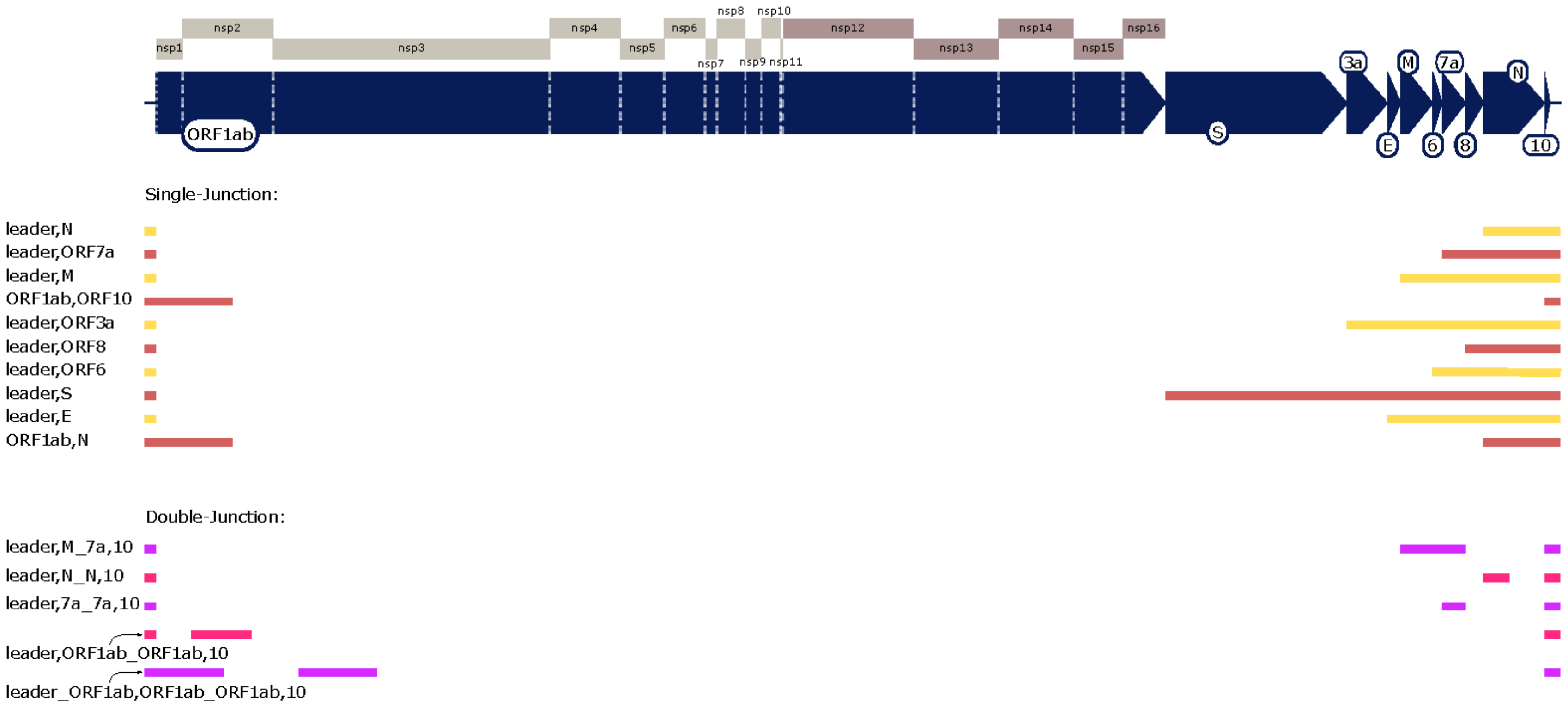

**Figure S1**: **Schematic of major sgRNA described in this study.** sgRNA are listed in order of abundance. Most of the central part of the genome is not expressed in subgenomic mRNA but is translated directly from genomic RNA.

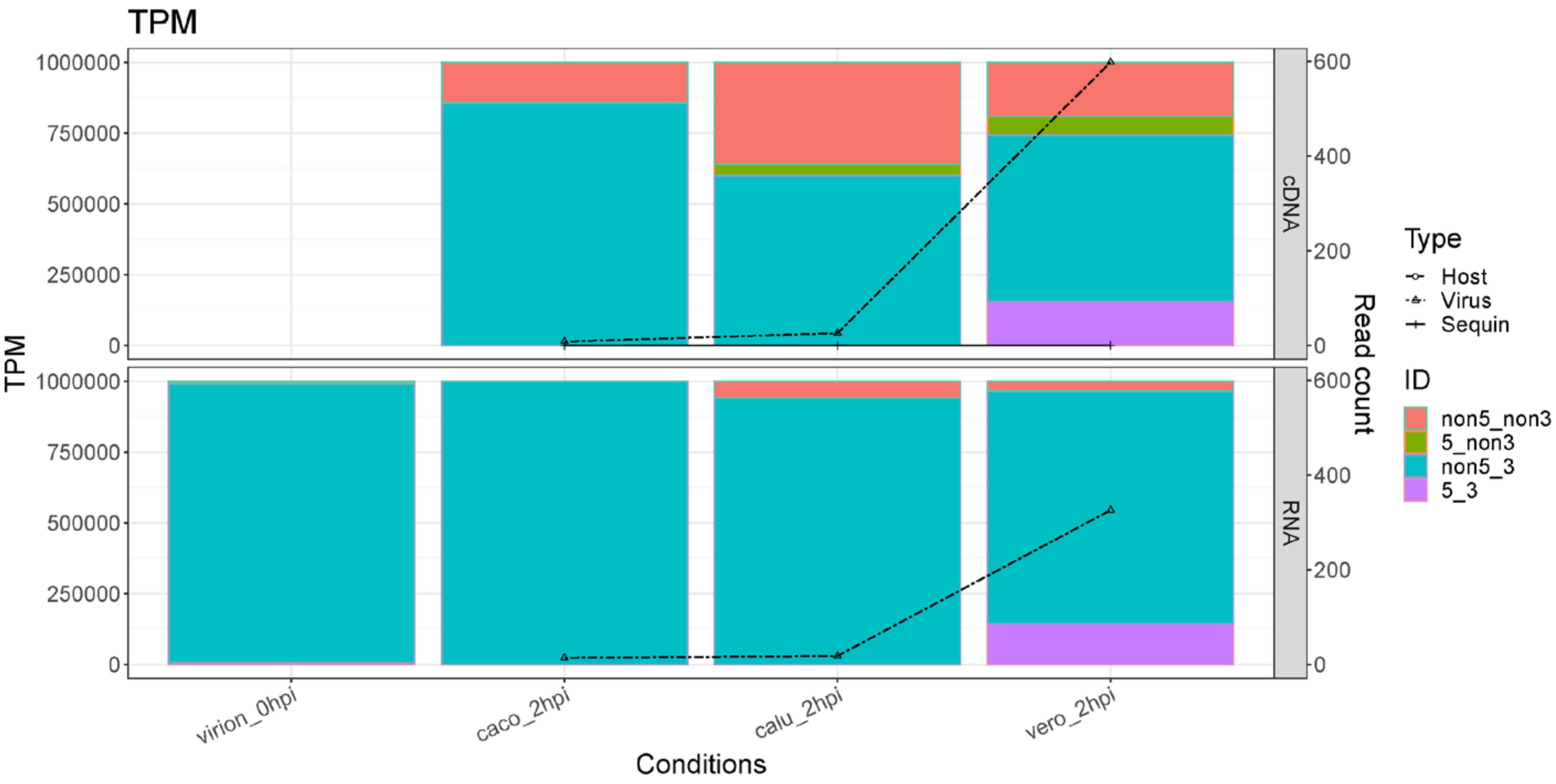

**Figure S2**: **Transcripts per million mapped viral reads for different read categories for RNA from infected cells 2 hpi (left axis) and total number of viral mapping reads (right axis)**. Top left cell is empty as virion cDNA was not sequenced. Related to **Figure 1**.

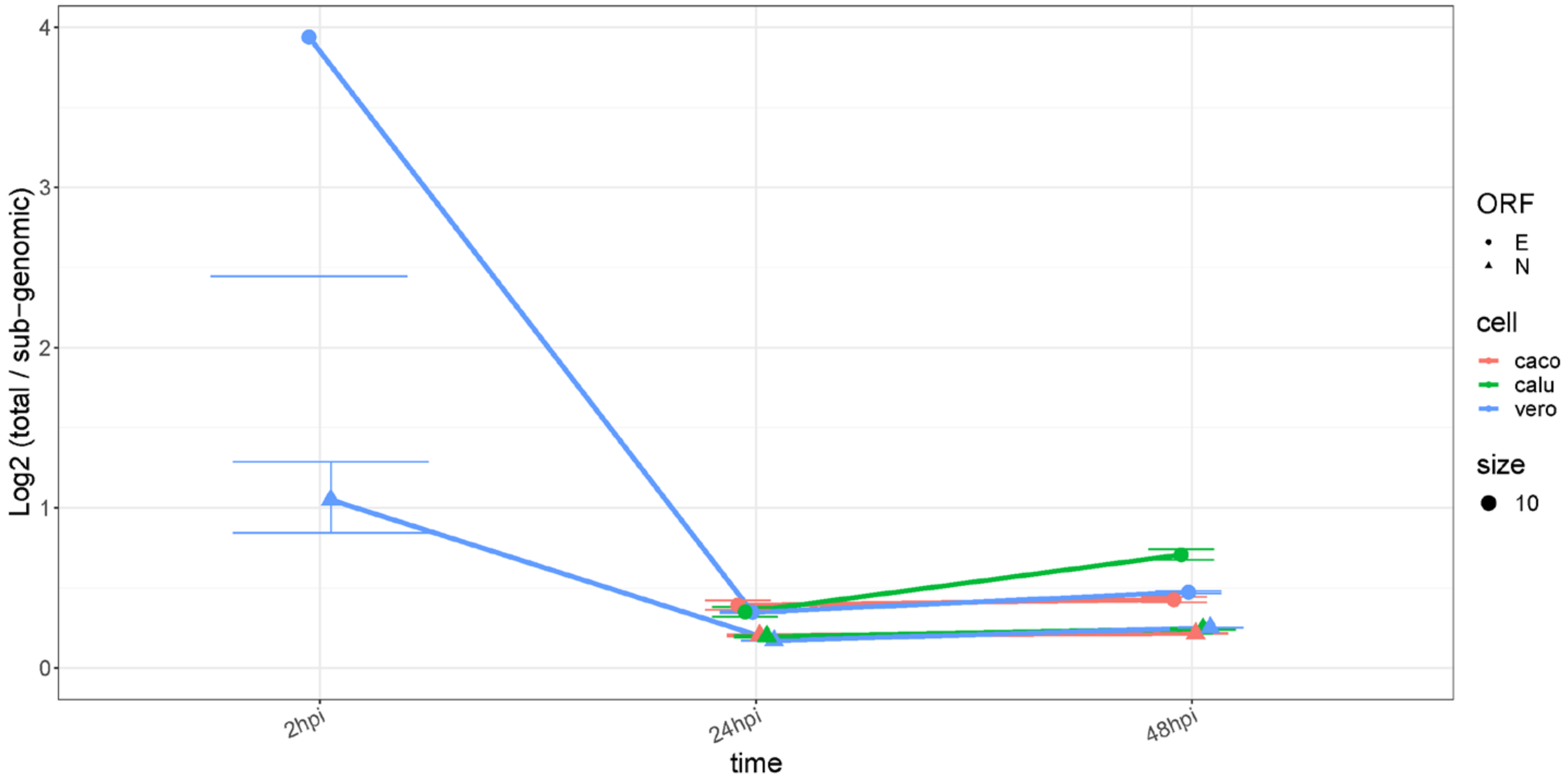

**Figure S3: Estimated Ct difference between sgRNA and total RNA from cDNA sequence data.** Error bars indicate 95% CI estimated based on read depth. For virion E 2 hpi, only the lower 95% CI is shown. Related to **Figure 1**.

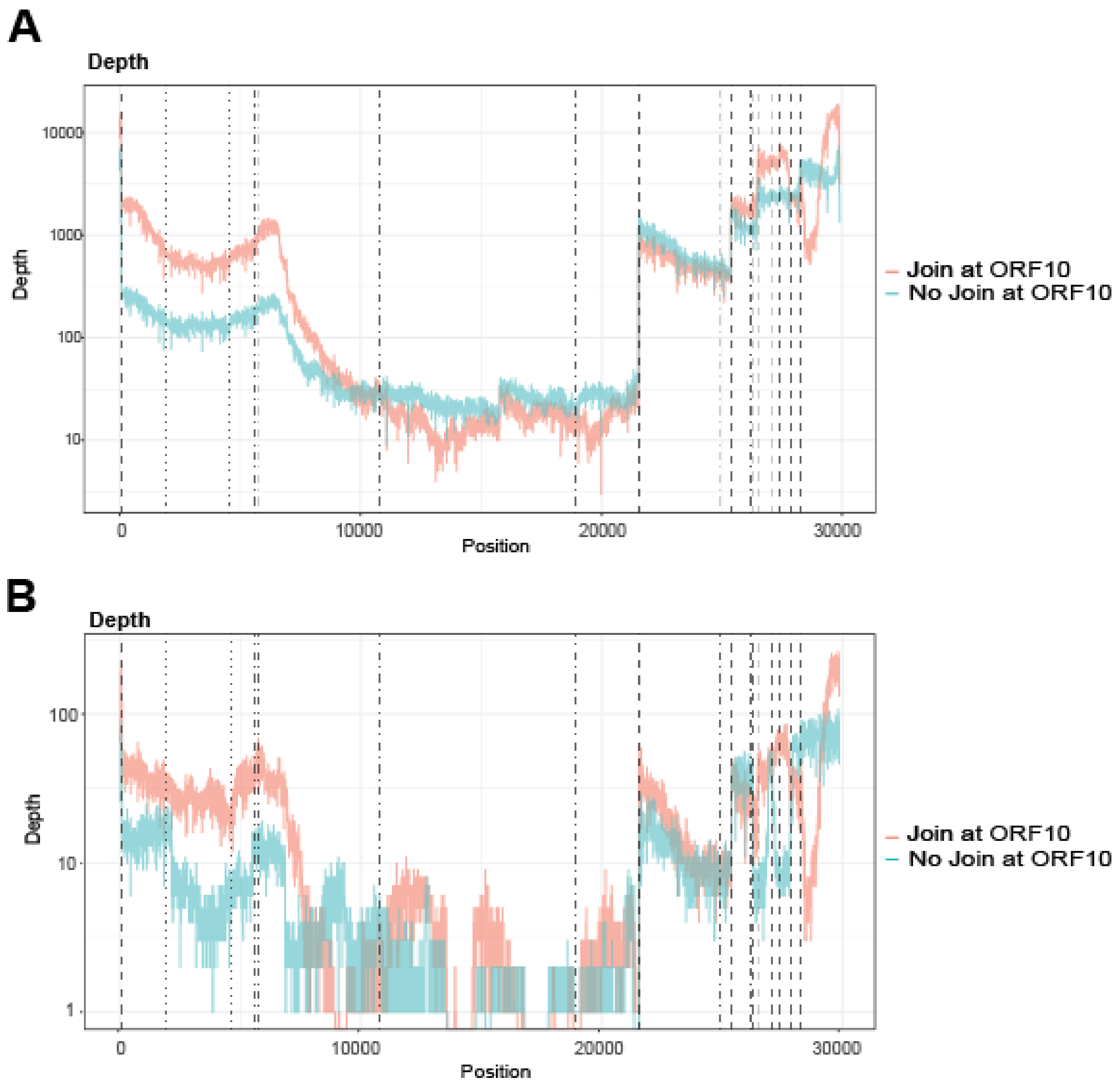

**Figure S4**: **Multiple junction transcript clusters. A.** Coverage of SARS-CoV-2 double-junction reads which have second breakpoint upstream of ORF10 (i.e. in the N ORF) vs those which do not. **B.** Coverage of SARS-CoV-2 triple junction reads which have second breakpoint upstream of ORF10 vs those which do not. Related to **Figure 4**.

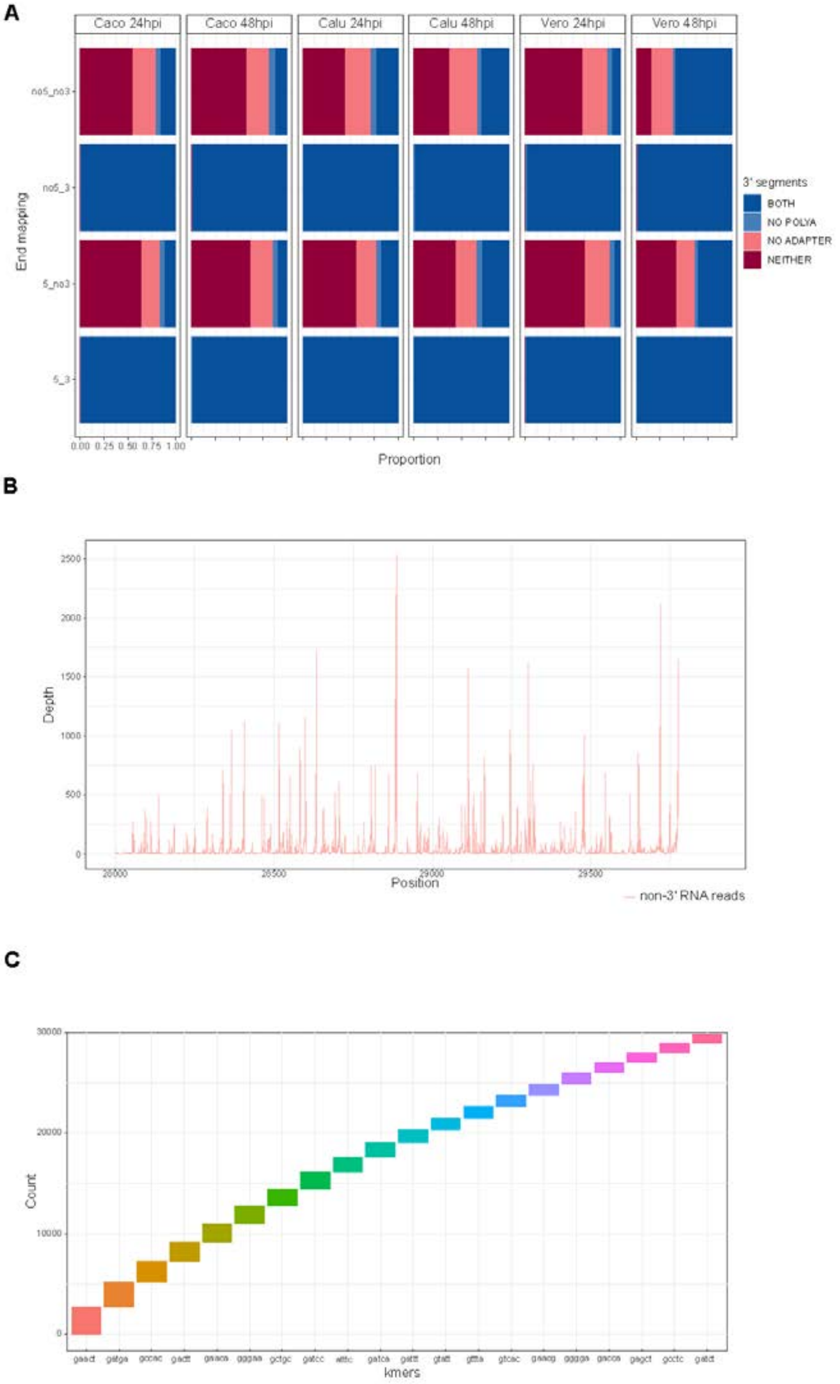

**Figure S5: Characterisation of non-3' mapping reads. A.** Poly(A) and sequencing adaptor detection from *nanopolish* 'polya'. Groups without 3' mapping from all experiments show consistent low detection of poly(A) tail. Vero 48 hpi has a comparatively high amount of poly(A) detection compared to other experiments which remains unexplained. **B.** Genome position at end of read for transcripts without 3' end within final 2000 bases of viral genome. There is non-random distribution of final breaks indicating unexplained bias in the reads that pass through the nanopore for sequencing. **C.** 5-mer at end of non-3' viral RNA reads. 5-mer is centered on final break of the transcript. Although non-random (**5B**), the sequencing of these transcripts cannot be explained by A-rich sequence at their ends. Related to **Table S1**.

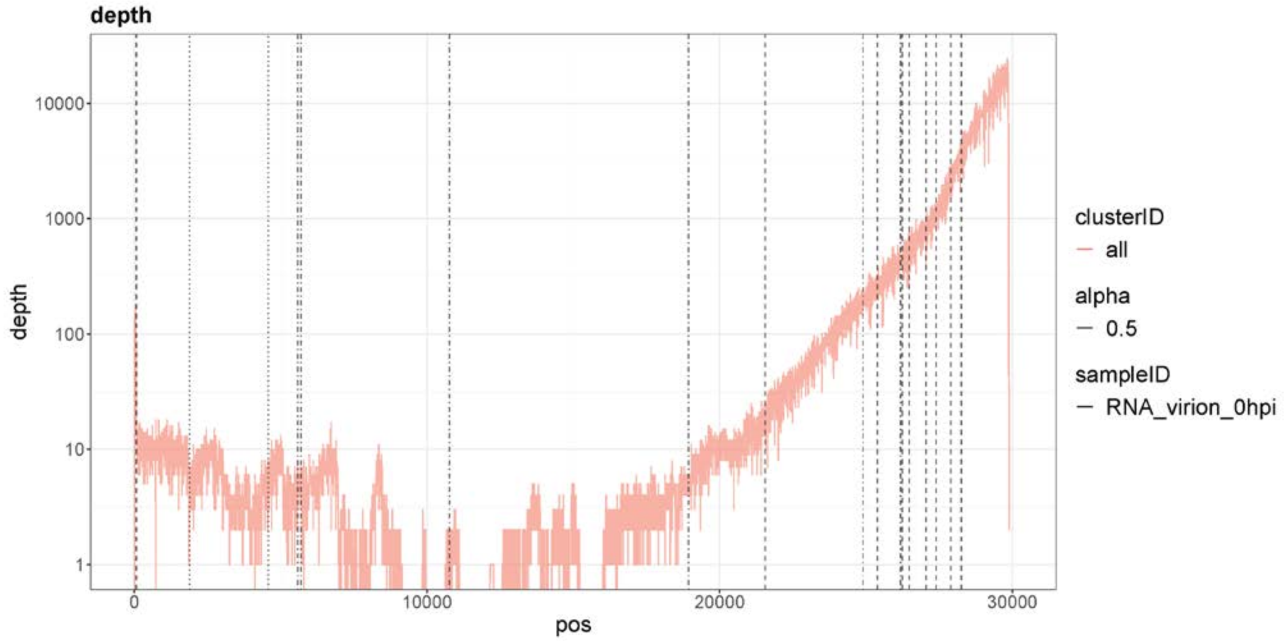

**Figure S6: Coverage of SARS-CoV-2 virion dataset.** Related to **Figure 6**.

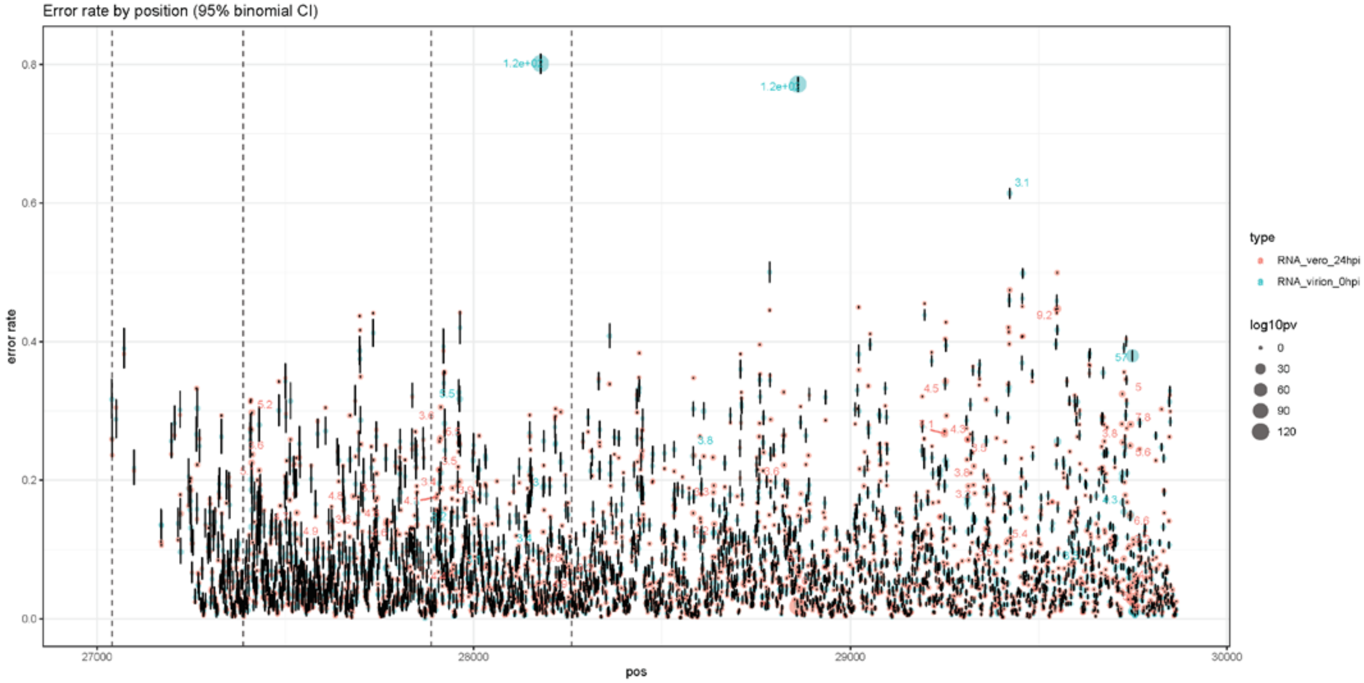

**Figure S7**: **Basecalling error rate by position for direct RNA extract from Virion (blue) and Vero at 24 hpi (orange).** Size of dot indicates the -$\log_{10}$ p-value of the fisher exact test comparing basecalling error between two conditions. Error bars indicate estimate 95% confidence interval of base-calling rate estimation (estimated using binomial model). Only sites for which both conditions have a coverage of at least 1000 are shown, and at all positions reads are sub-sampled down to a depth of 1000. Dashed vertical lines indicate position of TRS. Related to **Figure 6**.

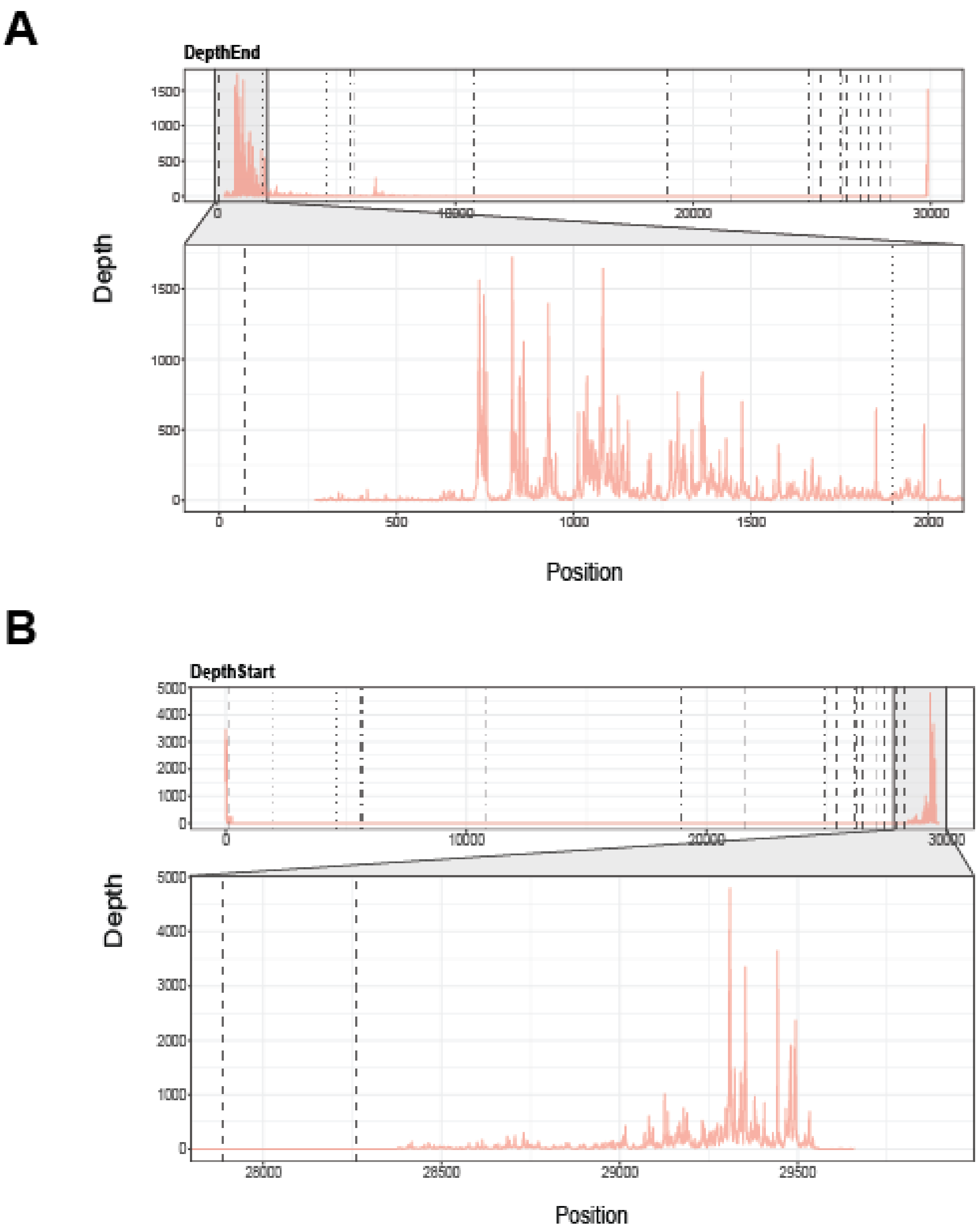

**Figure S8**: **Histogram of breakpoint positions for leader_ORF1ab,ORF10_3UTR transcript cluster. A.** Position at which reads assigned to cluster have a 5' breakpoint or terminate (i.e. contribute to depth "ending". Zoomed in region shows histogram of 5' break points up to 2 kb. **B.** Position at which reads assigned to cluster have 3' breakpoint or begin (i.e. contribute to depth "starting"). Zoomed in region shows histogram of 3' breakpoints in final 1.5 kb. Dashed vertical lines show position of TRS motif. Related to **Figure 5**.
